# Mixtures of large-scale dynamic functional brain network modes

**DOI:** 10.1101/2022.05.03.490453

**Authors:** Chetan Gohil, Evan Roberts, Ryan Timms, Alex Skates, Cameron Higgins, Andrew Quinn, Usama Pervaiz, Joost van Amersfoort, Pascal Notin, Yarin Gal, Stanislaw Adaszewski, Mark Woolrich

## Abstract

Accurate temporal modelling of functional brain networks is essential in the quest for understanding how such networks facilitate cognition. Researchers are beginning to adopt time-varying analyses for electrophysiological data that capture highly dynamic processes on the order of milliseconds. Typically, these approaches, such as clustering of functional connectivity profiles and Hidden Markov Modelling (HMM), assume mutual exclusivity of networks over time. Whilst a powerful constraint, this assumption may be compromising the ability of these approaches to describe the data effectively. Here, we propose a new generative model for functional connectivity as a time-varying linear mixture of spatially distributed statistical “modes”. The temporal evolution of this mixture is governed by a recurrent neural network, which enables the model to generate data with a rich temporal structure. We use a Bayesian framework known as amortised variational inference to learn model parameters from observed data. We call the approach DyNeMo (for Dynamic Network Modes), and show using simulations it outperforms the HMM when the assumption of mutual exclusivity is violated. In resting-state MEG, DyNeMo reveals a mixture of modes that activate on fast time scales of 100-150 ms, which is similar to state lifetimes found using an HMM. In task MEG data, DyNeMo finds modes with plausible, task-dependent evoked responses without any knowledge of the task timings. Overall, DyNeMo provides decompositions that are an approximate remapping of the HMM’s while showing improvements in overall explanatory power. However, the magnitude of the improvements suggests that the HMM’s assumption of mutual exclusivity can be reasonable in practice. Nonetheless, DyNeMo provides a flexible framework for implementing and assessing future modelling developments.

## 1 Introduction

Functional connectivity (FC, [1]) has traditionally been studied across the duration of an experiment, be it metabolic (e.g. [2, 3, 4, 5]) or electrophysiological in nature (e.g. [6, 7, 8, 9]). Such studies have shown that the brain forms well-defined spatio-temporal networks which are seen both in task [10] and at rest [11]. However, there is a growing body of evidence supporting the idea that these networks are transient [12, 13, 14], and that they emerge and dissolve on sub-second time scales. It is now well established that the dynamics of these networks underpin healthy brain activity and cognition [15] and that the disruption of FC is implicated in disease [16, 17].

A systematic understanding of the neuroscientific significance of these networks of wholebrain activity is only facilitated by accurate modelling across the spatial, temporal and spectral domains. Sliding window analyses have been used successfully to study time-varying FC in both M/EEG [18, 19, 20, 21, 22, 23, 24, 13, 25] and fMRI [26, 27, 28, 29, 30, 31, 32, 33, 34, 35]. Recent studies have calculated very short, or even instantaneous, time-point-by-time-point estimates of FC, which are then combined with a second stage of clustering such as k-means (e.g. [13]) to pool over recurrent patterns of otherwise poorly estimated FC. These two-stage approaches allow access to FC on fast time scales [36, 37].

Although they remain popular, sliding window analyses are a heuristic approach to data analysis and lack a generative model. An alternative approach to studying dynamics of functional brain networks is via the adoption of a formal model. An Hidden Markov Model (HMM) [38] is one such option. As with the two-stage approaches mentioned above, HMMs can pool non-contiguous periods of data together to make robust estimations of the activity of brain networks, including FC. However, they do so by incorporating these two stages into one model. HMMs (as well as other techniques, such as microstates [39]) have been used to show that brain networks evolve at faster time scales than previously suggested by competing techniques (such as independent component analysis) [14]. In the context of M/EEG, HMMs have been used to elucidate transient brain states [12], model sensor level fluctuations in covariance [40] and reveal latent task dynamics attributed to distributed brain regions [10]. More recently, Seedat et al. applied an HMM to detect transient bursting activity and showed it was correlated to aspects of the electrophysiological connectome [41], whilst Higgins et al. were able to show that replay in humans coincides with activation of the default mode network [42].

Although very powerful, convenient, and informative, traditional HMMs are themselves limited in two key ways. Firstly, there is the modelling choice that the state at any time point is only conditionally dependent on the state at the previous time point (i.e. the model is Markovian). This limits the modelling capability of the technique as there is no way for any long-range temporal dependencies between historic state occurrences and the current state to be established [43]. While approaches that use Hidden Semi-Markov Models have been proposed, they are limited in the complexity of long-range temporal dependencies they can capture [44]. Secondly, HMMs adopt a mutually exclusive state model, meaning that data can only be generated by one set of observation model parameters at any given instance. True brain dynamics might be better modelled by patterns that can flexibly combine and mix over time. The mutual exclusivity constraint was found to lead to errors in inferred functional brain network metrics in [45].

We set to address these two limitations in this paper and do so by introducing a new generative model for neuroimaging data. Specifically, we model the time-varying mean and covariance of the data as a linear weighted sum of spatially distributed patterns of activity or “modes”. Notably, we do not impose mutual exclusivity on mode activation. Similarly, we drop the assumption that the dynamics of the modes are a function of a Markovian process. This is achieved by using a unidirectional recurrent neural network (RNN) [46] to model the temporal evolution of the weighted sum. The memory provided by the RNN facilitates a richer context to the changes in the instantaneous mean and covariance than what would be afforded by a traditional HMM.

In this work, we use Bayesian methods [47] to infer the parameters of the generative model. With this method, we learn a distribution for each parameter, which allows us to incorporate uncertainty into our parameter estimates. Having observed data, we update the distributions to find likely parameters for the model to have generated the data. In this work, we adapt a method used in variational autoencoders [48] to infer the model parameter distributions. One component of this is amortised inference, which works through the deployment of an inference network. In our case the inference network is another RNN, which is bidirectional [46] and learns a mapping from the observed data to the model parameter distributions. The use of an inference network facilitates the scaling and application of this technique to very large datasets, without ever needing (necessarily) to increase the number of inference network parameters to be learnt.

To update our model parameter distributions, we minimise the variational free energy (see Section 2.2) using stochastic gradient descent [46]. We do this by sampling from the model parameter distributions using the reparameterisation trick [48]. The ability to estimate the variational free energy by sampling enables us to use sophisticated generative models that include highly non-linear transformations that would not be feasible with classical Bayesian methods. Taken together, we call the generative model and inference framework DyNeMo (Dynamic Network Modes).

## 2 Methods

In this section we outline the generative model and describe the inference of model parameters. We also describe the datasets and preprocessing steps carried out in this work.

### 2.1 Generative Model

Here we propose a model for generating neuroimaging data that explicitly models functional brain networks, including a metric of their FC, as a dynamic quantity. The model describes time series data using a set of *modes*, which are constituent elements that can be combined to define time-varying statistics of the data. When trained on neuroimaging data, modes are simply static spatial brain activity patterns that can overlap with each other. We refer to them as “modes” to emphasise that the model is not categorical, i.e. that modes should not be mistaken for mutually exclusive *states* (as would be the case in an HMM). Similar to an HMM, our generative model has two components: a latent representation and a data generating process given the latent representation, which is referred to as an *observation model*. In our case, the latent representation is a set of mixing coefficients ***α***_*t*_ and the observation model is a multivariate normal distribution. The mean and covariance of the multivariate normal distribution is determined by linearly mixing the modes’ spatial models, i.e. means ***μ***_*j*_ and covariances ***D***_*j*_, with the coefficients ***α***_*t*_. The mixing coefficients are dynamic in nature whereas the modes are static. Therefore, dynamics in the observed data are captured in the dynamics of the mixing coefficients. The mixing coefficients provide a low-dimensional and interpretable dynamic description of the data and modes correspond to static spatial distributions of activity/FC, where mode-specific FC is captured by the between-brain-region correlations in ***D***_*j*_. Both of these quantities are useful for understanding the data. An overview of the generative model is shown in Figure 1 and a mathematical formulation is given below.

**Figure 1:**
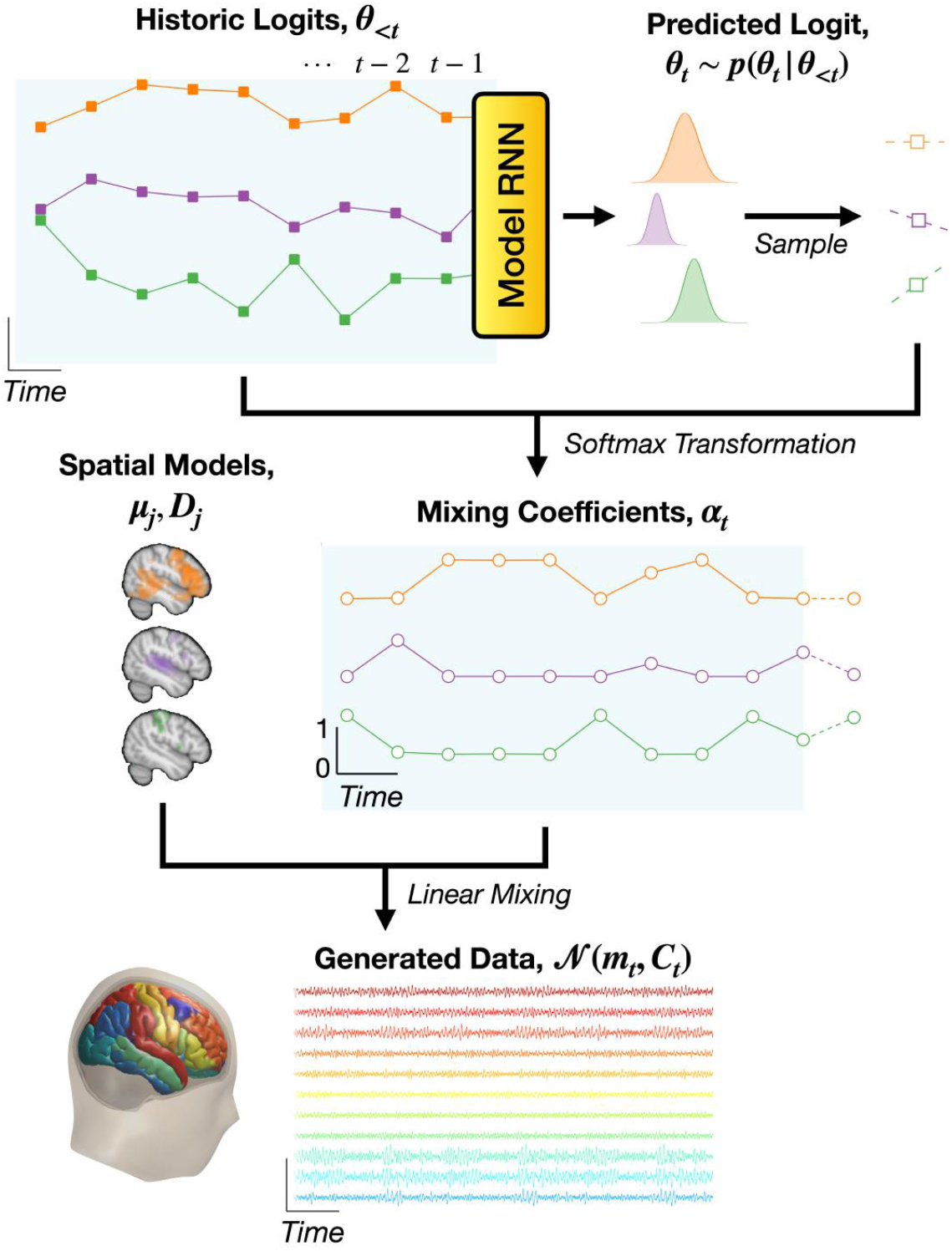
Generative model employed in DyNeMo. Historic values of a latent logit time series (solid squares, blue background), ***θ***_<*t*_, are fed into a unidirectional model RNN. The output of the model RNN parameterises a normal distribution, *p*(***θ***_*t*_|***θ***_<*t*_), which we sample to predict the next logit, ***θ***_*t*_, (unfilled squares). These logits are transformed via a softmax operation to give the mixing coefficients, ***α***_*t*_, (unfilled circles). The softmax transformation enforces the mixing coefficients are positive and sum to one at any instance in time. Separate from the dynamics are the corresponding spatial models that describe brain network activity as a set of modes (depicted in different colours here); via a mean vector, and covariance matrix, ***D***_*j*_. The mode spatial models combine with the dynamic mixing coefficients (linear mixing) to parameterise a multivariate normal distribution with a time-varying mean vector, ***μ***_*t*_, and covariance matrix, ***C***_*t*_. Note, we do not enforce any constraint on the modes means ***μ***_*j*_ and covariances ***D***_*j*_, this means they can overlap in time and space and the overall activity (***m***_*t*_ and ***C***_*t*_) can vary.

At each time point *t* there is a probabilistic vector of free parameters, referred to as a *logit* and denoted by ***θ***_*t*_. The logits are distributed in accordance with a multivariate normal distribution,

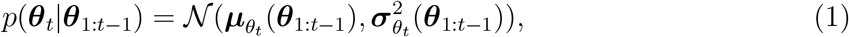

where ***θ***_1:*t*–1_ denotes a sequence of historic logits {***θ***_1_,…, ***θ***_*t*–1_}, ***μ***_*θ*_*t*__ is a mean vector and 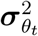 is a diagonal covariance matrix. We use a unidirectional RNN to predict future values of ***μ***_*θ*_*t*__ and ***σ***_*θ*_*t*__ based on previous logits ***θ***_1:*t*–1_. The logit at each time point ***θ***_*t*_ is sampled from the distribution *p*(***θ***_*t*_|***θ***_1:*t*–1_). The historic values of the logits ***θ***_1:*t*•1_ are fed into the RNN:

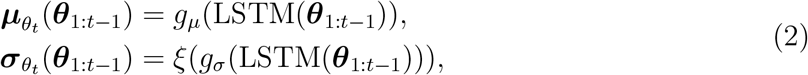

where *g_μ_* and *g_σ_* are learnt affine transformations, is a softplus function included to ensure the standard deviations ***σ***_*θ*_*t*__ are positive, and LSTM is a type of RNN known as a Long Short Term Memory network [49]. We refer to this network as the *model RNN*. The logits ***θ***_*t*_ are used to determine a set of mixing coefficients,

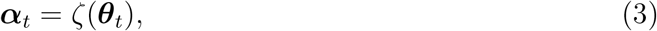

where *ζ* is a softmax function which assures that the ***α***_*t*_ values are positive and sum to one.^1^ The mixing coefficients are then used together with a set of spatial modes to calculate a time-varying mean vector and covariance matrix:

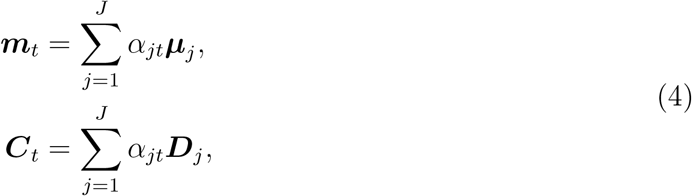

where *J* is the number of modes, ***μ***_*j*_ is the mean vector for each mode, ***D***_*j*_ is the covariance matrix for each mode and *α*_*jt*_ are the elements of ***α***_*t*_.

### 2.2 Inference

In this section we describe the framework employed to infer the parameters of our generative model. Namely, the logits ***θ***_*t*_, mode means ***μ***_*j*_ and covariances ***D***_*j*_. In this work, we use variational Bayesian inference to learn the full posterior distribution for ***θ***_*t*_ and point estimates for ***μ***_*j*_ and ***D***_*j*_.

### Variational Bayes

In Bayesian inference, we would like to learn a distribution, referred to as the *posterior distribution*, for the variable we are trying to estimate given some data we have observed. In *variational* Bayesian inference, we approximate the posterior distribution with a simple distribution, referred to as the *variational posterior distribution q*(***θ***_*t*_), and aim to minimise the Kullback-Leibler (KL) divergence between the variational and true posterior, which amounts to minimising the variational free energy (or equivalently, maximising the evidence lower bound). In classical variational Bayes [51, 52, 53], this involves formulating update rules for the parameters of the variational posterior distribution given some observed data. Deriving these update rules is only made possible by limiting the complexity of the generative model for the observed data and restricting the variational posterior to conjugate distributions. In addition to this, we have a separate variational distribution for each variable we are trying to estimate. Also in classical variational Bayes, we learn the parameters of each variational distribution separately, which becomes problematic in terms of computer memory requirements when we wish to estimate a large number of variables.

In brief, we overcome these difficulties with a technique adapted from variational autoencoders [48]. This deploys a neural network (which we call the *inference network*) to perform amortised inference, which helps the approach to scale to large numbers of observations over time; and a sampling technique (known as the *reparameterisation trick*) that allows us to learn a full posterior distribution for ***θ***_*t*_ [48]. We learn point estimates of ***μ***_*j*_ and ***D***_*j*_ using trainable free parameters. We update estimates for ***μ***_*j*_, ***D***_*j*_, and the posterior distribution parameters of ***θ***_*t*_, to minimise the variational free energy using stochastic gradient descent.

### Logits *θ*_*t*_

Focusing on the full posterior inference of the logits ***θ***_*t*_, here, we use amortised inference [52]. This involves using an inference network to learn a mapping from the observed data to the parameters of the variational posterior. The rationale for this approach is that the computation from past inferences can be reused in future inferences. The use of an inference network fixes the number of trainable parameters to the number of internal weights and biases in the inference network. This is usually significantly smaller than the number of time points, which allows us to efficiently scale to bigger datasets.

### Inference network

We now describe the inference network in detail. Having observed the time series ***x***_1:*N*_, we approximate the variational posterior distribution for ***θ***_*t*_ as

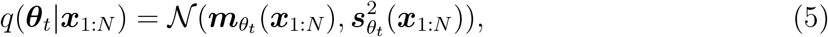

where ***m***_*θ_t_*_ and 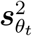 are the variational posterior mean and covariance of a multivariate normal distribution respectively. The variational posterior covariance is a diagonal matrix. We use a bidirectional RNN for the inference network, which we refer to as the *inference RNN*. This network outputs the parameters of the variational posterior distribution given the observed data:

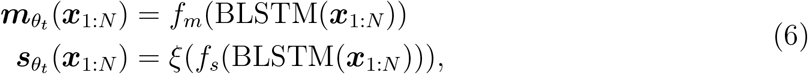

where *f_m_* and *f_s_* are affine transformations and BLSTM denotes a bidirectional LSTM. The complete DyNeMo framework and interplay between the generative model and inference network is shown in Figure 2.

**Figure 2:**
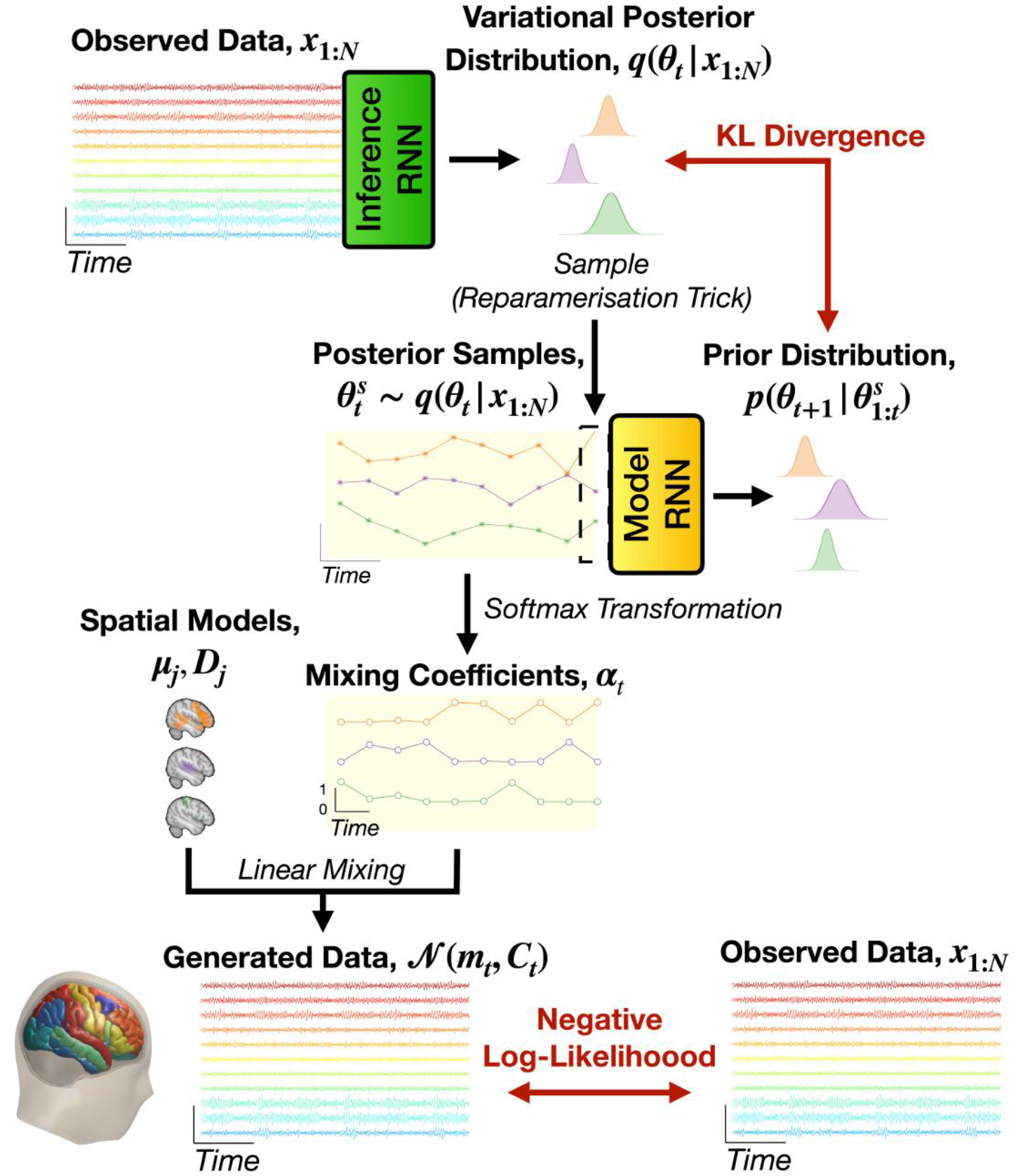
The full DyNeMo framework. A sequence of observed data, ***x***_1:*N*_, is fed into a bidirectional RNN which parameterises the approximate variational posterior distribution for the logit time series, *q*(***θ***_*t*_|***x***_1:*N*_). We sample 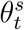 from the variational posterior distribution using the reparameterisation trick (asterisks, orange background) and feed the samples into the model RNN to predict the prior distribution one time step in the future *p*(***θ***_*t*+1_ |***θ***_1:*t*_). The prior and posterior distribution are used to calculate the KL divergence term of the variational free energy. The samples from the variational posterior distribution 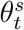 are also used to generate the observed data by first applying a softmax transformation to calculate the mixing coefficients, ***α***_*t*_, (unfilled circles, orange background). These mixing coefficients are then combined with the spatial model of each mode, which is a mean vector, ***μ***_*j*_, and covariance matrix, ***D***_*j*_. This gives an estimate of the time-varying mean, ***m***_*t*_, and covariance, ***D***_*j*_, which is used to calculate the negative log-likelihood term of the variational free energy.

### Loss function

Having outlined the inference network for the logits, we turn our attention to the loss function used in DyNeMo. In variational Bayesian inference we infer a parameter, in this case ***θ***_*t*_, by minimising the variational free energy [54],

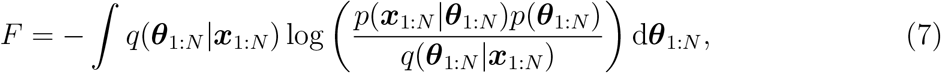

where *p*(***θ***_1:*N*_) is the prior and *p*(***x***_1:*N*_|***θ***_1:*N*_) is the likelihood. With this approach the inference problem is cast as an optimisation problem, which can be efficiently solved with the use of stochastic gradient descent [46]. Here, we make stochastic estimates of a loss function, and use the gradient of the loss function to update the trainable parameters in our model. However, to estimate the loss function we must calculate the integral in Equation (7). In DyNeMo, this is done using a sampling technique (i.e. the reparameterisation trick) to give Monte Carlo estimates of the loss function.

Insight into the loss function is gained by re-writing Equation (7) as two terms (see SI 9.1):

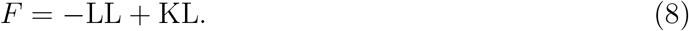

The first term is referred to as the *log-likelihood term* and the second term is referred to as the *KL divergence term.* The log-likelihood term acts to give the most probable estimate for the logits that could generate the training data and the KL divergence term acts to regularise the estimate. Relating this to components of DyNeMo, it is the inference RNN that infers the logits, which together with the learnt mode means and covariances determine the loglikelihood term, whilst the model RNN regularises the inferred logits through its role as the prior in the KL divergence term. It is the temporal regularisation provided by the model RNN that distinguishes DyNeMo from a Gaussian mixture model (GMM). The benefit of including a model RNN for temporal regularisation is discussed in SI 9.4.

We now detail the calculation used to estimate the loss function. The log-likelihood term is given by

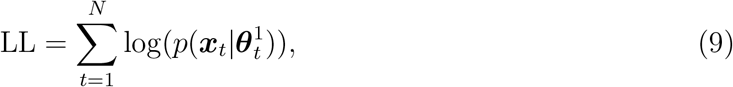

where 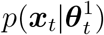 is the likelihood of generating data ***x***_*t*_ at time point *t* given the latent variable is 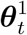, which is a sample from the variational posterior *q*(***θ***_*t*_|***x***_1:*N*_). The superscript in 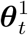 indicates that it is the first sample from *q*(***θ***_*t*_|***x***_1:*N*_). Only one sample from the variational posterior at each time point is used to estimate the log-likelihood term. Note that the likelihood is a multivariate normal whose mean and covariance is determined by Equation (4). Therefore, the likelihood depends on the logits ***θ***_*t*_, mode means ***μ***_*j*_ and covariances ***D***_*j*_. The KL divergence term is given by

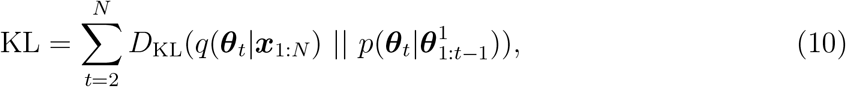

where 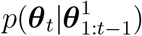 is the prior distribution for ***θ***_*t*_ given a single sample for the previous logits 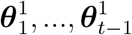 from their respective variational posteriors *q*(***θ***_1_|***x***_1:*N*_),…, *q*(***θ***_*t*-1_|***x***_1:*N*_) and *D*_KL_ is the KL divergence [51] between the variational posterior and prior. A full derivation of the loss function is given in SI 9.1.

### Reparameterisation trick

Next, we outline the method used to sample from the variational posterior distribution **q**(***θ***_*t*_|***x***_1:*N*_). This is a multivariate normal distribution with mean vector ***m***_*θ*_*t*__(***x***_1:*N*_) and diagonal covariance matrix 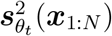. To obtain a sample 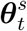 from *q*(***θ***_*t*_|***x***_1:*N*_), we use the reparameterisation trick [48], where we sample from a normal distribution,

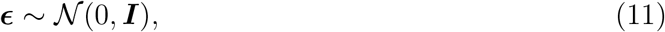

where ***I*** is the identity matrix. ***ϵ***^*s*^ denotes the *s*^th^ sample from 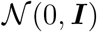. We calculate the samples for the logits as

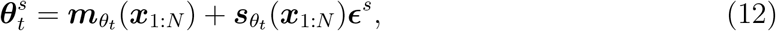

where ***s***_*θ*_*t*__(***x***_1:*N*_) is a vector containing the square root of the diagonal from 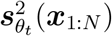. The use of the reparameterisation trick allows us to directly minimise the loss function using stochastic gradient descent.

### Mode means *μ_j_* and covariances *D_j_*

Having detailed the inference of the logits ***θ***_*t*_ and the calculation of the loss function, we now turn our attention to the spatial models described by the means ***μ***_*j*_ and covariances ***D***_*j*_. We performed fully Bayesian inference on the logits, as they are temporally local parameters, and hence will have reasonably large amounts of uncertainty in their estimation which needs to be propagated to the inference of ***θ***_*t*_ over time. By contrast, the mode means ***μ***_*j*_ and covariances ***D***_*j*_ are global parameters whose inference can draw on information over all time points. As a result we choose to use point estimates for ***μ***_*j*_ and ***D***_*j*_, which are learnt using trainable free parameters. Additionally, learning point estimates when they are sufficient has the advantage of simplifying inference.

The time-varying mean vector ***m***_*t*_ constructed from the mode means ***μ***_*j*_ can take on any value, and can therefore be treated as free parameters. However, the time-varying covariance ***C***_*t*_ constructed from the ***D***_*j*_ matrices is required to be positive definite. We enforce this by parameterising the ***D***_*j*_’s using the Cholesky decomposition,

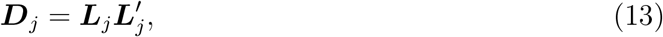

where ***L***_*j*_ is a lower triangular matrix known as a Cholesky factor and’ denotes the matrix transpose. We learn ***L***_*j*_ as a vector of free parameters that is used to fill a lower triangular matrix. We also apply a softplus operation and add a small positive value to the diagonal of the Cholesky factor to improve training stability. Using this approach, we learn point estimates for the mode means and covariances.

### Hyperparameters, initialisation and training

The full DyNeMo model contains several hyperparameters, for example the number of layers and hidden units in the RNNs, the batch size, the learning rate, and many more. These all must be specified before training the model. DyNeMo also contains a large number of trainable parameters, which must be initialised. A description of the hyperparameters and the initialisation of trainable parameters is given in SI 9.2. Hyperparameters for each dataset used in this work are summarised in Table 1. There are also several techniques that can be used to improve model training, such as KL annealing [55] and using multiple starts. These are also discussed in detail in SI 9.2.

**Table 1:** Hyperparameters (see SI 9.2) used in simulation and real data studies.

| Hyperparameter | Simulation 1 | Simulation 2 | MEG Data |
| --- | --- | --- | --- |
| Number of modes, $J$ | 3 | 6 | 10 |
| Sequence length, $N$ | 200 | 200 | 200 |
| Inference RNN hidden units | 64 | 64 | 64 |
| Model RNN hidden units | 64 | 64 | 64 |
| KL annealing sharpness, $A_S$ | 10 | 10 | 10 |
| KL annealing epochs, $n_{AE}$ | 100 | 100 | 300 |
| Training epochs $n_E$ | 200 | 200 | 600 |
| Batch size | 16 | 16 | 32 |
| Learning rate, $\eta$ | 0.01 | 0.01 | 0.0025 |
| Gradient clip (norm.) | - | - | 0.5 |
| Number of multi-starts | - | - | 10 |
| Multi-start epochs | - | - | 20 |

## 2.3 Datasets

In this section, we describe the data used to train DyNeMo. This includes simulated data, described in Sections 2.3.1 and 2.3.2, which was used to evaluate DyNeMo’s modelling and inference capabilities, and real MEG data, described in Section 2.3.3, which was used for neuroscientific studies.

### 2.3.1 Simulation 1: Long-Range Dependencies

The first simulation dataset was used to examine DyNeMo’s ability to learn long-range temporal dependencies in the underlying logits. In simulation 1, data were generated using a Hidden Semi-Markov Model (HSMM) [56]. Unlike an HMM, state lifetimes are explicitly modelled in an HSMM. This enables us to specify a lifetime distribution where long-lived states are probable. We train DyNeMo on this data and examine samples from the generative model, in this case we sample the model RNN. The lifetime distribution of the sampled states indicates the memory of the model RNN, i.e. the time scale of temporal dependencies it has learnt. If samples from DyNeMo show long-lived states that cannot be generated with an HMM, we say DyNeMo has learnt long-range temporal dependencies. In simulation 1, we used a gamma distribution (with shape and scale parameters of 5 and 10 respectively) to sample state lifetimes. We use a transition probability matrix with self-transitions excluded to determine the sequence of states to sample a lifetime for. The transition probability matrix and ground truth mode covariances are shown in Figures 4a and 4b respectively. A multivariate time series with 11 channels, 25,600 samples and 3 hidden states was generated using an HSMM simulation with a multivariate normal observation model. A zero mean vector was used for each mode and covariances were generated randomly. The ground truth state time course and lifetime distribution of this simulation is shown in Figures 4c and 4d respectively.

### 2.3.2 Simulation 2: Linear Mode Mixing

The second simulation dataset was used to examine DyNeMo’s ability to infer a linear mixture of co-activating modes. Here, we simulated a set of *J* sine waves with different amplitudes, frequencies and initial phases to represent the logits ***θ***_*t*_. We applied a softmax operation at each time point to calculate the ground truth mixing coefficients ***α***_*t*_. A multivariate normal distribution with zero mean and randomly generated covariances was used for the observation model. A multivariate time series with 80 channels, 25,600 samples and 6 hidden modes was simulated. The first 2,000 time points of the simulated logits and mixing coefficients are shown in Figures 5a and 5b respectively.

### 2.3.3 MEG Data

In addition to the simulation datasets, we trained DyNeMo on two real MEG datasets: a resting-state and a (visuomotor) task dataset. The MEG datasets were source reconstructed to 42 regions of interest. The raw data, preprocessing and source reconstruction are described below.

#### Raw data and preprocessing

Data from the UK MEG Partnership were used in this study. The data were acquired using a 275-channel CTF MEG system operating in third-order synthetic gradiometry at a sampling frequency of 1.2 kHz. Structural MRI scans were acquired with a Phillips Achieva 7 T. MEG data were preprocessed using the OHBA software library (OSL, [57]). The time series was downsampled to 250 Hz before a notch filter at 50 Hz (and harmonics) was used to remove power line noise. The data were then bandpass filtered between 1 and 98 Hz. Finally, an automated bad segment detection algorithm in OSL was used to remove particularly noisy segments of the recording. No independent component analysis was applied to identify artefacts.

#### Source reconstruction

Structural data were coregistered with the MEG data using an iterative close-point algorithm; digitised head points acquired with a Polhemous pen were matched to individual subject’s scalp surfaces extracted with FSL’s BET tool [58, 59]. We used the local spheres head model in this work [60]. Preprocessed sensor data were source reconstructed onto an 8 mm isotropic grid using a linearly constrained minimum variance beamformer [61]. Voxels were then parcellated into 42 anatomically defined regions of interest, before a time series for each parcel was extracted by applying Principal Component Analysis (PCA) to each region of interest. We use the same 42 regions of interest as [62], see the supplementary information of [62] for a list of the regions used and their MNI coordinates. Source reconstruction can lead to artefactual correlations between parcel time courses, referred to as *source leakage*. This is a static effect so it should not affect the inference of dynamics. However, it can affect the inferred FC. We minimise source leakage using the symmetric multivariate leakage reduction technique described in [63], which unlike pairwise methods has the benefit of reducing leakage caused by so-called ghost interactions [64]. We will refer to each parcel as a *channel*.

#### Resting-state dataset

The resting-state dataset is formed from the MEG recordings of 55 healthy participants (mean age 38.3 years, maximum age 62 years, minimum age 19 years, 27 males, 50 right handed). The participants were asked to sit in the scanner with their eyes open while 10 minutes of data were recorded.

#### Task dataset

The task dataset is formed from MEG recordings of 51 healthy participants (mean age 38.4 years, maximum age 62 years, 24 males, 46 right handed). The recordings were taken while the participants performed a visuomotor task [65]. Participants were presented with a high-contrast grating (visual stimulus). The grating remained on screen for a jittered duration between 1.5 and 2 seconds. When the grating was removed, the participants performed an abduction using the index finger and thumb of the right hand. This abduction response was measured using an electromyograph on the back of the hand. From the grating removal, an 8 second inter trial interval is incorporated until the grating re-appeared on the screen. The structure of the task is shown in Figure 3. A total of 1,837 trials are contained in this dataset. The majority of participants in the UK MEG Partnership study have both resting-state and task recordings. 48 of the participants in the resting-state and task dataset are the same.

**Figure 3:**
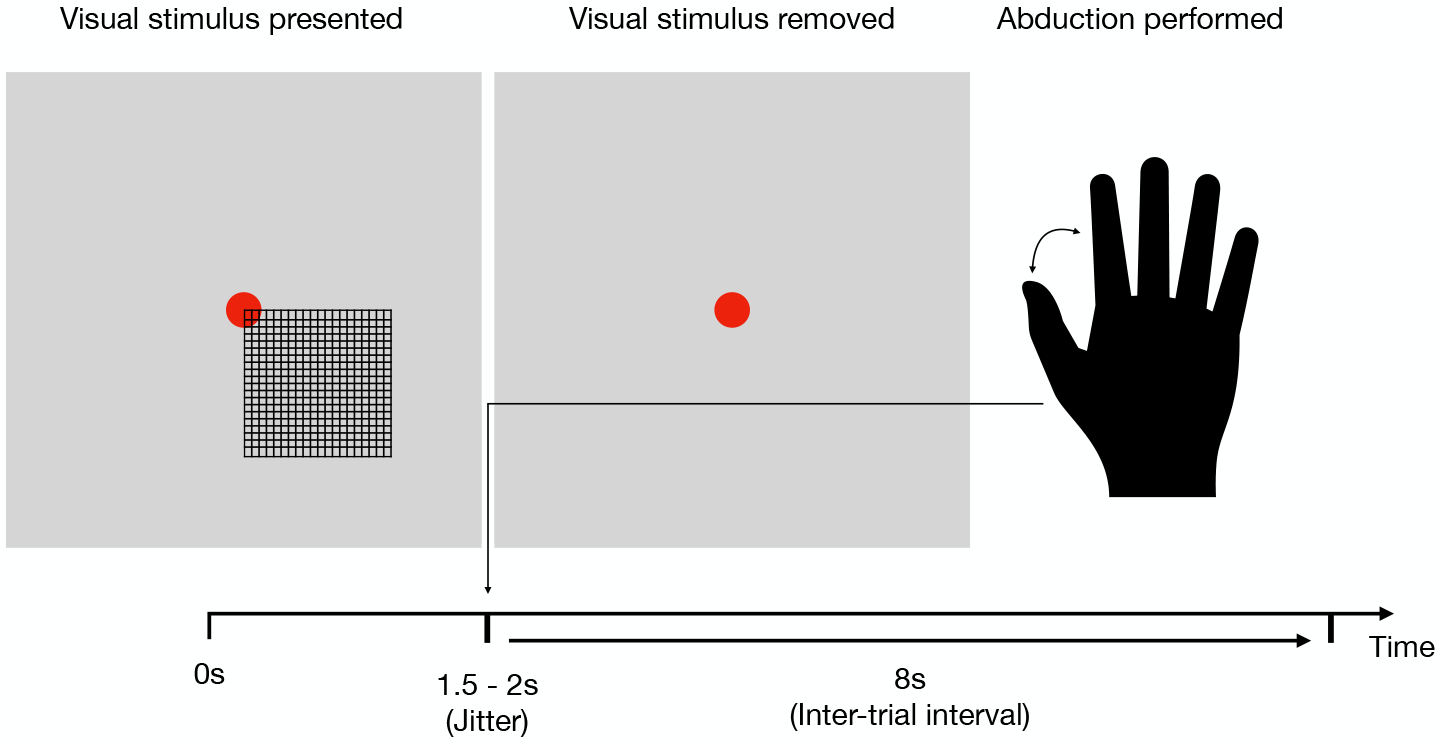
The structure of the visuomotor task. Participants are presented with a visual stimulus, which is an onscreen grid. After a period of between 1.5 and 2 seconds, the grid is removed. Upon grid removal, the participant performs a right-hand index finger abduction. Between the removal of the grid and its reappearance for the next trial, there is an 8 second inter-trial interval.

#### Data preparation

Before training DyNeMo, we further prepare the preprocessed data by performing the following steps. The first step is used to encode spectral information into the observation model (see Figure S1), whereas the other two are to help train the model. These steps are optional and were only performed on the MEG datasets. The steps are:

1. Time-delay embedding. This involves adding extra channels with time-lagged versions of the original data. We use 15 embeddings, which results in a total of 630 channels. By doing this, we introduce additional off-diagonal elements to the covariance matrix, which contains the covariance of a channel with a time-lagged version of itself. This element of the covariance matrix is the autocorrelation function of the channel for a given lag [66]. As the autocorrelation function captures the spectral properties of a signal, this allows the model to learn spectral features of the data as part of the covariance matrix.
2. PCA. After time-delay embedding we are left with 630 channels. This is too much for modern GPUs to hold in memory. Therefore, we use PCA for dimensionality reduction down to 80 channels.
3. Standardisation (z-transform) across the time dimension. This is a common transformation that has been found to be essential in many optimisation problems [46]. Standardisation is the final step in preparing the training data.^2^

Time-delay embedding and PCA are summarised in Figure S1. We train DyNeMo to generate the prepared MEG data, i.e. the 80 channel time series after time-delay embedding and PCA, rather than the 42 channel time series of source reconstructed data.

#### 2.4 Post-hoc Analysis of Learnt Latent Variables

In this work, we set each mode’s mean vector, ***μ***_*j*_, to zero and do not update its value during training. This is due to our choice of training data. In the simulation datasets, we simulated modes with a zero mean vector so there is no need to model the mean. In the MEG datasets, we train on time-delay embedded data. Here, we want all the spectral information to be contained in the mode covariance matrices, therefore we set the means to zero. Additionally, we would like to compare our results to those presented in [62], which trained an HMM without learning the mean. In this work, we use DyNeMo to learn the mixing coefficients, ***α***_*t*_, (via the logits, ***θ***_*t*_) and the mode covariances, ***D***_*j*_.

DyNeMo provides a variational posterior distribution *q*(***θ***_*t*_|***x***_1:*N*_) at each time point. To simplify analysis we take the most probable value for ***θ***_*t*_ (this is known as the *maximum a posteriori probability estimate*) and use this to calculate the inferred mode mixing coefficients, ***α***_*t*_, which contain a description of latent dynamics in the training data.^3^

We can use the inferred mode mixing coefficients to estimate quantities that characterise the training data. We describe such analyses in detail in SI 9.3. Quantities calculated in the post-hoc analyses include: summary statistics that characterise the temporal properties of each mode, such as activation lifetimes, interval times and fractional occupancies; power spectra that characterise the spectral properties of each mode and power/FC maps that characterise the spatial pattern of each mode. Note, we only use the the inferred mixing coefficients (and the source reconstructed data) in the post-hoc analysis, the mode covariances are not used.

## 3 Results

### 3.1 Simulation 1: Long-Range Dependencies

A simulation dataset was used to examine DyNeMo’s ability to learn long-range temporal dependencies. DyNeMo was trained on the simulation dataset described in Section 2.3.1. An HMM was also trained on the simulated data for comparison. In this simulation, a mutually exclusive hidden state was used to generate the training data. The ground truth hidden state time course is shown in Figure 4c. DyNeMo was able to correctly infer mutually exclusive modes, which we can think of as states. The DyNeMo and HMM inferred state time courses are also shown in Figure 4c. Both DyNeMo and the HMM are able to infer the presence of long-range dependencies by matching the ground truth, non-exponential, state lifetime distributions (shown in Figure 4d). A dice coefficient (model inferred vs ground truth) of greater than 0.99 is achieved for both models. However, this does not mean that the HMM or DyNeMo generative models have necessarily learnt long-range dependencies, as the inferred state time courses could be a result of purely data-driven information. To test this, we can sample state time courses from the trained HMM and DyNeMo generative models and examine their lifetime distributions. Figure 4e shows the lifetime distribution sampled state time courses. The state lifetime distribution of the sample from DyNeMo captures the nonexponential ground truth distribution, demonstrating its ability to learn long-range temporal dependencies over the scale of at least 50 samples. Contrastingly, the HMM was not able to generate any long-range temporal dependencies, indicating that, as expected, it is only able to capture short-range dependencies.

**Figure 4:**
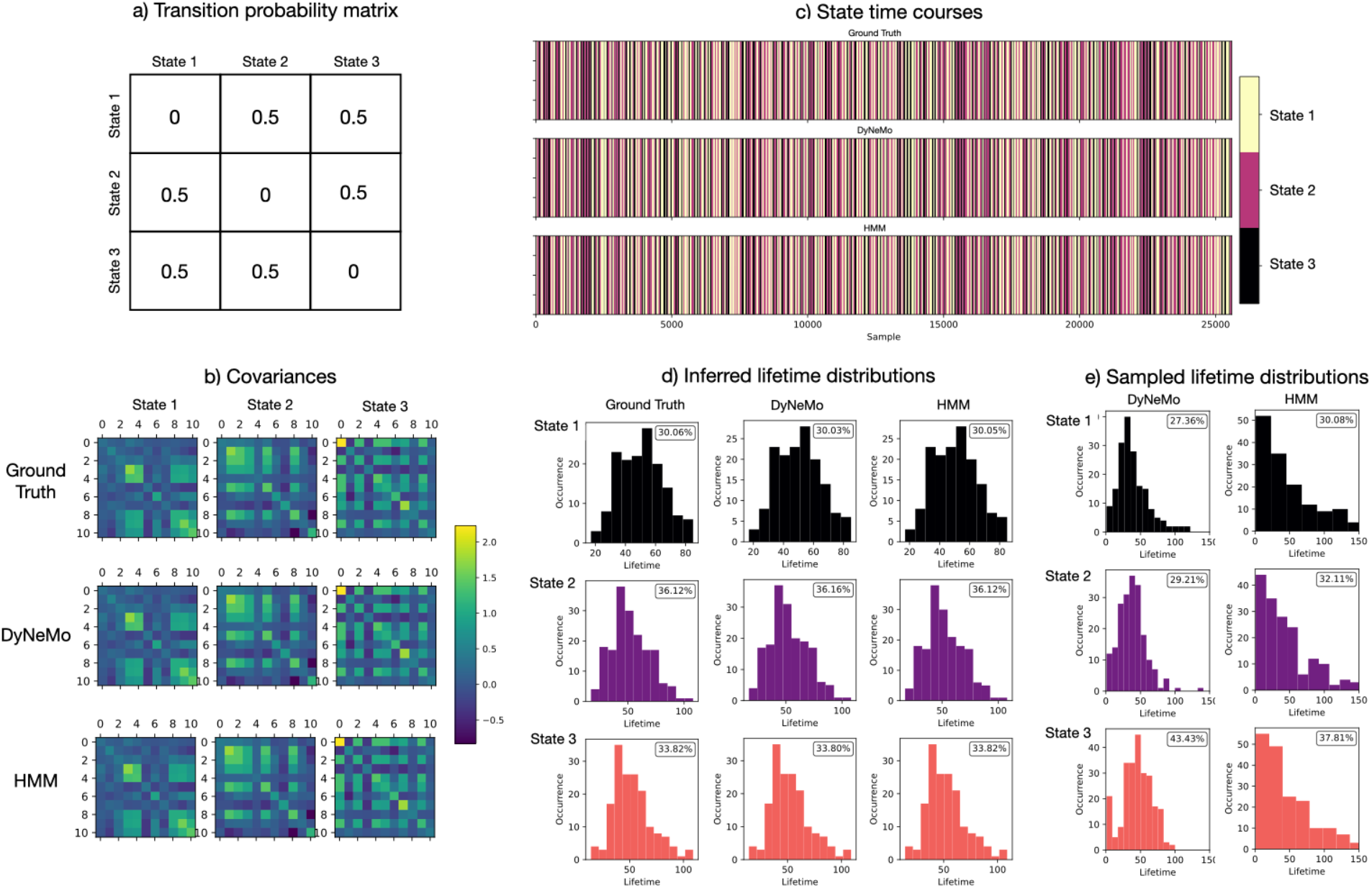
DyNeMo is able to learn long-range temporal dependencies in the latent dynamics of simulated data. Parameters of an HSMM simulation are shown along with the parameters inferred by DyNeMo and an HMM. While both DyNeMo and the HMM were able to accurately *infer* the hidden state time course and their lifetime distributions, actual *samples from each model* show that only DyNeMo was able to learn the lifetime distribution of the states within its generative model, demonstrating its ability to learn long-range temporal dependencies. a) Transition probability matrix used in the simulation. b) Covariances: simulated (top), inferred by DyNeMo (middle) and inferred by an HMM (bottom). c) State time courses: simulated (top), inferred by DyNeMo (middle) and inferred by an HMM (bottom). Each colour corresponds to a separate state. d) Lifetime distribution of inferred state time courses. e) Lifetime distribution of sampled state time courses. The fractional occupancy of each state is shown as a percentage in each histogram plot.

### 3.2 Simulation 2: Linear Mode Mixing

In contrast to the mutual exclusivity assumption of the HMM, DyNeMo has the ability to infer a linear a mixture of modes. To test this we trained DyNeMo and the HMM for comparison on the simulation dataset described in Section 2.3.2. Figure 5b shows the simulated mixing coefficients and those inferred by DyNeMo. For comparison, the state time course inferred by an HMM is also shown in Figure 5c. As the HMM is a mutually exclusive state model, it is unable to infer a linear mixture of modes, whereas DyNeMo’s mixing coefficients estimate the ground truth very well, demonstrating its ability to learn a mixture of modes. Using the inferred mixing coefficients or state time course along with the inferred covariances, we can reconstruct the time-varying covariance, ***C***_*t*_, of the training data. The Riemannian distance between the reconstruction and ground truth is shown in Figure 5d. The mean Riemannian distance for DyNeMo is 1.5, whereas it is 11.9 for the HMM. Using a paired t-test the difference is significant with a *p*-value < 10^-5^. The smaller Riemannian distance indicates DyNeMo is a more accurate model for the time-varying covariance.

**Figure 5:**
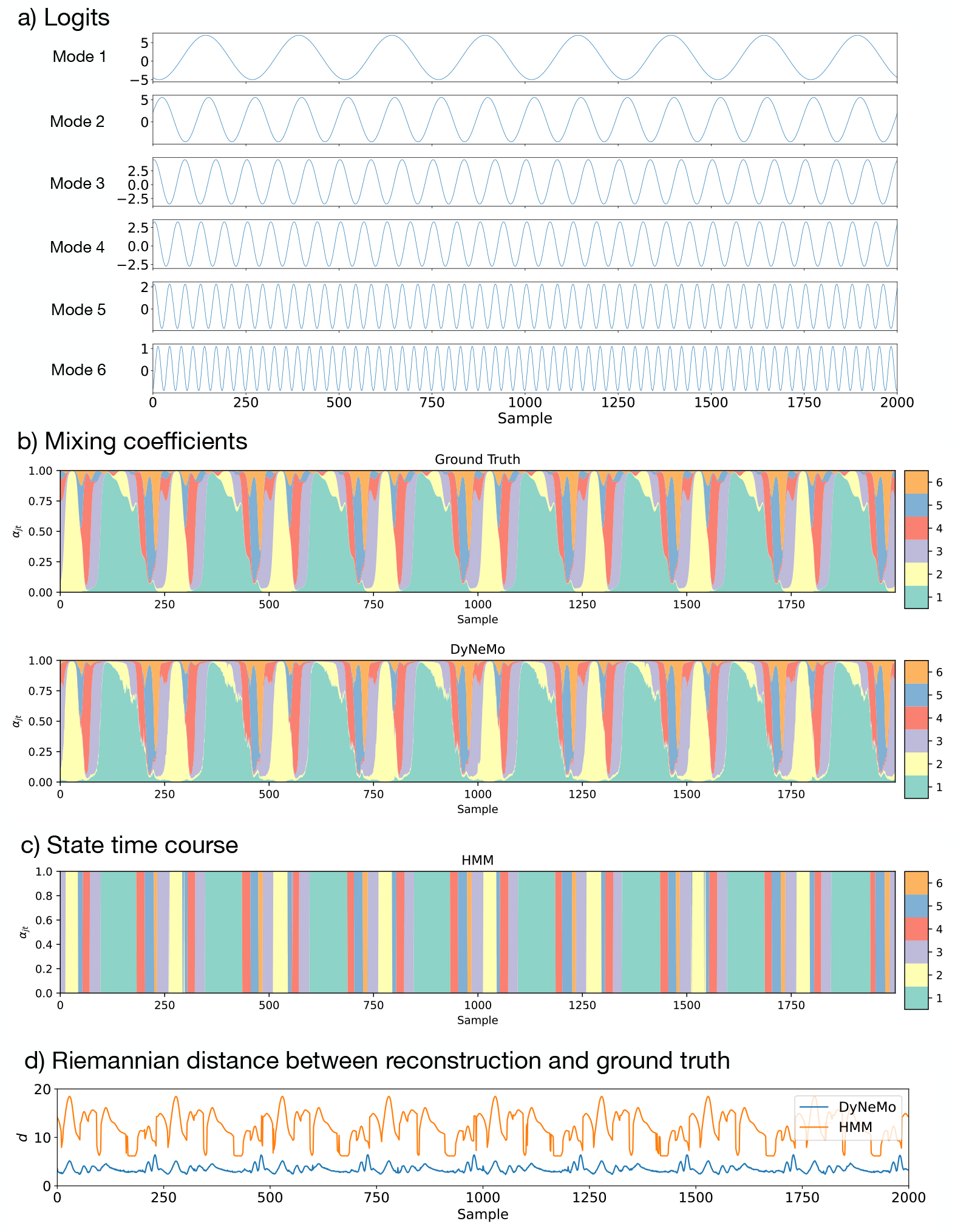
DyNeMo is able to accurately infer a linear mixture of modes. DyNeMo was trained on a simulation with co-activating modes. The mixing coefficients inferred by DyNeMo follow the same pattern as the ground truth. The failure of an HMM in modelling this type of simulation due to its inherent assumption of mutual exclusivity is also shown. a) Logits used to simulate the training data. b) Mixing coefficients of the simulation (top) and inferred by DyNeMo (bottom). c) State time course inferred by an HMM. d) Riemannian distance between the reconstruction of the time-varying covariance, ***C***_*t*_, (via Equation (4)) and the ground truth for DyNeMo and the HMM. Only the first 2000 time points are shown in each plot.

### 3.3 Resting-State MEG Data

#### DyNeMo identifies plausible resting-state networks

Figure 6 shows the power maps, FC maps and power spectral densities (PSDs) of 10 modes inferred by DyNeMo when trained on the resting-state MEG dataset described in Section 2.3.3. For the PSDs, we plot the regression coefficients ***P***_*j*_ (*f*) to highlight differences relative to the mean PSD ***P***_0_(*f*) common to all modes. Mode 1 appears to be a low-power background network and does not show any large deviations in power from the mean PSD for any frequency. Modes 2-10 show high power localised to specific regions associated with functional activity (see [67] for an overview of the functional association of different brain networks). Regions with high power also appear to have high FC. Modes 2 and 3 show power in regions associated with visual activity. Mode 4 shows power in parietal regions and can be associated with the posterior default mode network (see Figure 11). Mode 5 shows power in the sensorimotor region. Modes 6-8 show power in auditory/language regions. Modes 2-8 show power in the alpha band (8-12 Hz) and modes 4-6 and 8 include power at higher frequencies in the beta band (15-30 Hz). Mode 9 shows power in fronto-parietal regions and is recognised as an executive control network. Mode 10 shows power in frontal regions which can be associated with the anterior default mode network. Modes 9 and 10 exhibit low-frequency oscillations in the delta/theta band (1-7 Hz). The PSD of each mode is consistent with the expected oscillations at the high-power regions in each mode [68]. A comparison with states inferred with this dataset using an HMM is presented in the section “Large-scale resting-state networks can be formed from a linear mixture of modes”.

**Figure 6:**
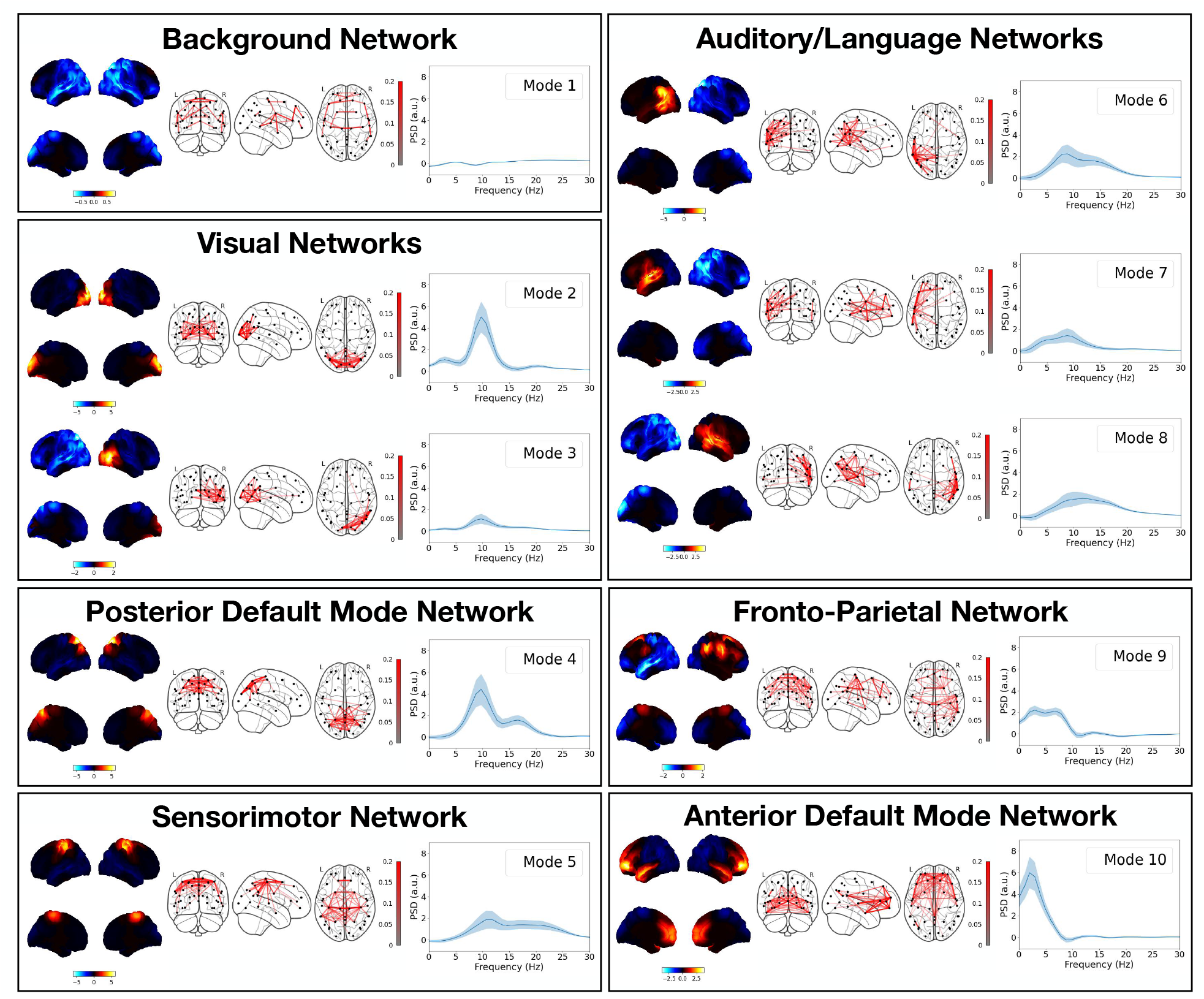
DyNeMo infers modes that form plausible resting-state MEG networks. Ten modes were inferred using resting-state MEG data from 55 subjects. Mode 1 appears to be a low-power background network, whereas modes 2-10 show high power in areas associated with functional networks. Modes are grouped in terms of their functional role. Each box shows the power map (left), FC map (middle) and PSD relative to the mean averaged over regions of interest (right) for each group. The top two views on the brain in the power map plots are lateral surfaces and the bottom two are medial surfaces. The shaded area in the PSD plots shows the standard error on the mean.

#### Power maps are reproducible across two split-halves of the dataset

To assess the reproducibility of modes across datasets, we split the full dataset into two halves of 27 subjects. We assess the reproducibility of the modes across halves using the RV coefficient [69], which is a generalisation of the squared Pearson correlation coefficient. We match the modes across halves in a pairwise fashion using the RV coefficient as a measure of similarity. Figure 7 shows the power maps of the matched modes. In general, the same regions are active in each pair of modes and the functional networks are reproducible across datasets. The main difference is small changes in how power is distributed across the visual network modes (mode 4) and across the temporal/frontal regions (mode 9).

**Figure 7:**
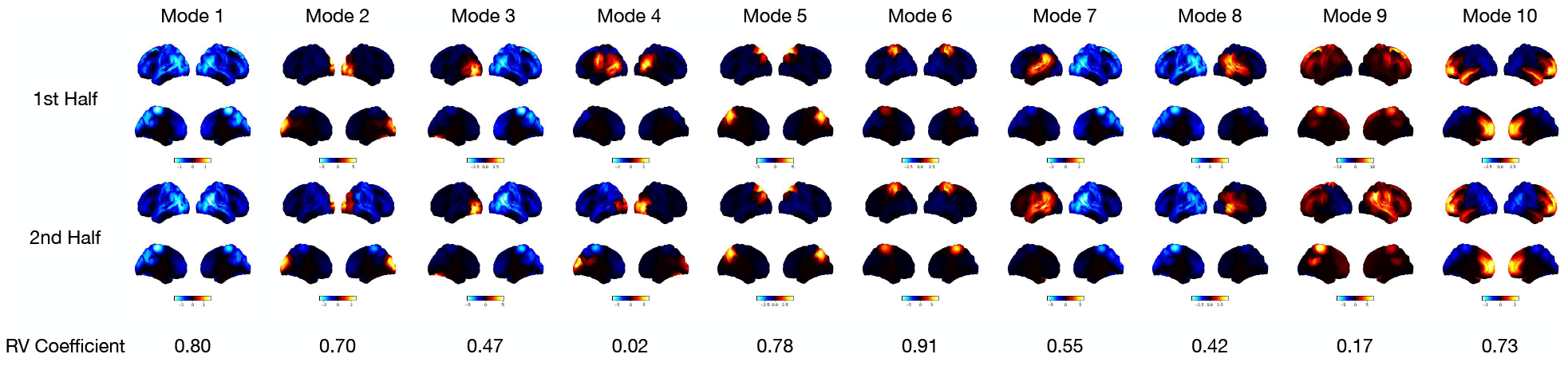
Power maps are reproducible across two split-halves of a dataset. Each half of the dataset contains the resting-state MEG data of 27 subjects. Power maps are shown for the the first half of the dataset (top) and second half of the dataset (middle). The RV coefficient of the inferred covariances from each half for a given mode (bottom) is also shown. The modes were matched in terms of their RV coefficient. Pairing the modes from each half we see the same functional networks are inferred. These networks also match the modes inferred on the full dataset of 55 subjects, suggesting these networks are reproducible across datasets. The top two views on the brain in each power map plot are lateral surfaces and the bottom two are medial surfaces.

#### Mode activations are anti-correlated with a background mode and modes with activity in similar regions co-activate

A subset of the inferred mixing coefficients is shown in Figure 8. Figure 8a shows the raw mixing coefficients inferred directly from DyNeMo. However, these mixing coefficients do not account for a difference in the relative magnitude of each mode covariance. For example, a mode with a small mixing coefficient may still be a large contributor to the time-varying covariance if the magnitude of its mode covariance is large. We can account for this by obtaining a weighted mixing coefficient mode time course by multiplying the raw mixing coefficients with the trace of its mode covariance. We also normalise the weighted mixing coefficient time course by dividing by the sum over all modes at each time point to maintain the sum-to-one constraint. Figure 8b and 8c show these normalised weighted mixing coefficients. Once we account for the magnitude of the mode covariances, we see each mode’s contribution to the time-varying covariance is roughly equal. We show the state time course inferred by an HMM in Figure 8d for comparison. Figure 8e shows the correlation between the raw mixing coefficients *αj_t_* for each mode. Modes 2-10 appear to be anti-correlated with mode 1. This arises due to the softmax operation (Equation (36)) that constrains the mixing coefficients to sum to one. For a mode to activate by contributing more to the time-varying covariance, another mode’s contribution must decrease. The anti-correlation of mode 1 with every other mode suggests that it is primarily this mode’s contribution that is decreased. This suggests that mode 1 can be thought of as a background mode that is deactivated by the other modes.

**Figure 8:**
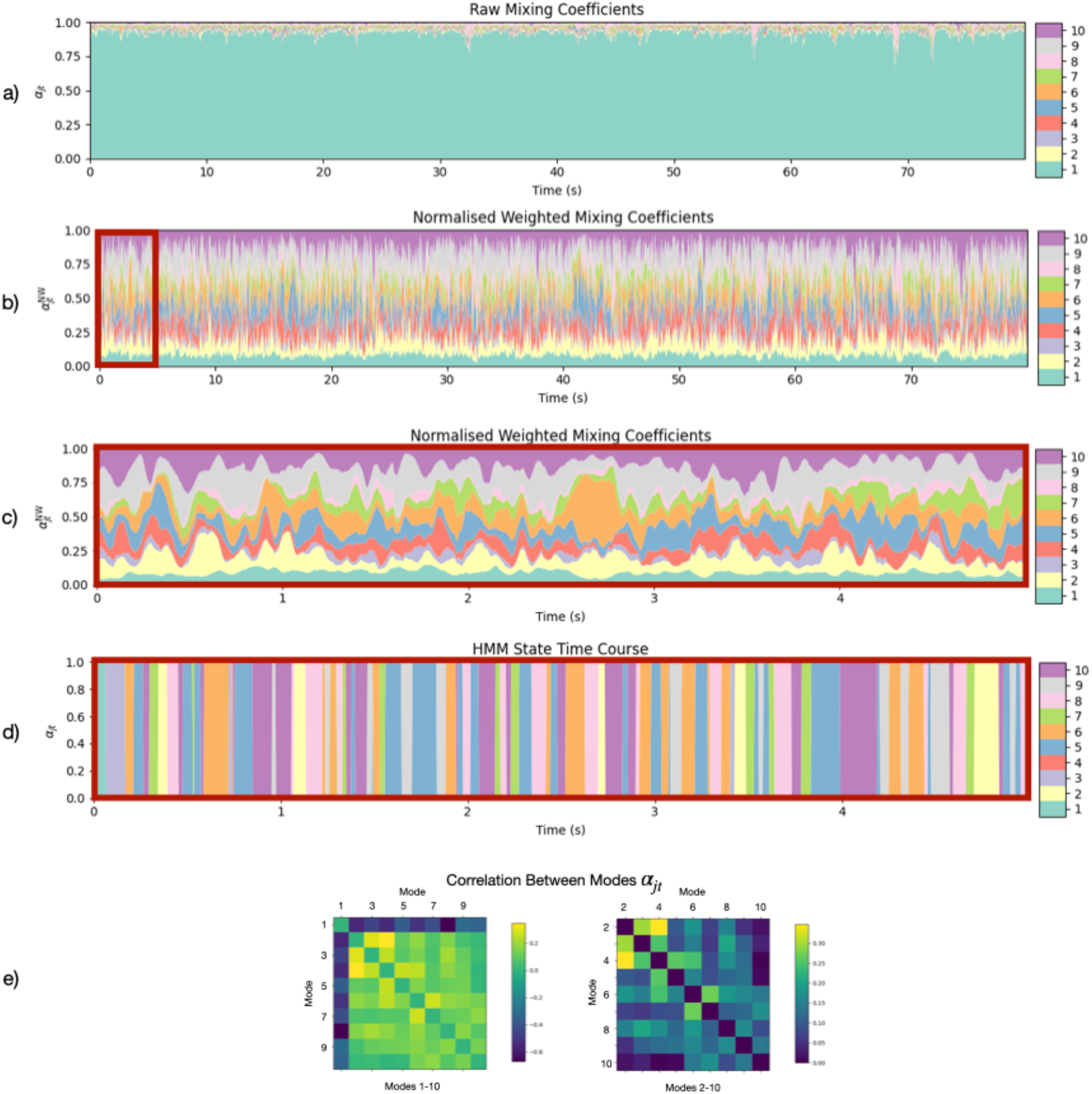
DyNeMo provides a mode description of resting-state MEG data. a) Raw mixing coefficients *α_jt_* inferred by DyNeMo for one subject. b) Mixing coefficients *α_jt_* weighted by the trace of each mode covariance and normalised to sum to one at each time point. c) Zoomed in normalised weighted mixing coefficients 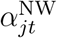 for the first 5 seconds. d) HMM state time course for the first 5 seconds for comparison. The power/FC maps and PSDs for the HMM states are shown in Figure S7. e) Correlation between the raw mixing coefficients *α_jt_* for different modes *j*. Ordering is the same as Figure 6. We see DyNeMo’s description of the data is a set of co-existing modes whose contribution to the time-varying covariance fluctuates. Once weighted by the covariance matrices we see each mode has a more equal contribution. We also see modes 2-10 are anti-correlated with the mode 1 and modes with activation in similar regions, e.g. modes 2, 3 and 4, are correlated.

#### DyNeMo reveals short-lived (100-150 ms) mode activations

Using a GMM to define when a mode is active we calculate summary statistics such as lifetimes, intervals and fractional occupancies. Mode activation time courses and summary statistics are shown in Figure 9. Mode 1 appears to have long activation lifetimes and a high fractional occupancy, which is consistent with the description of it being a background network that is largely present throughout. Modes 2-10 have mean lifetimes approximately over the range 100-150 ms, which is slightly longer than the state lifetimes obtained from an HMM, which are over the range 50-100 ms [62]. Both models reveal transient networks with lifetimes on the order of 100 ms, suggesting that this is a plausible time scale for these functional networks in resting-state MEG data, confirming that the short lifetimes previously found by the HMM are not likely to be caused by the mutual exclusivity assumption.

**Figure 9:**
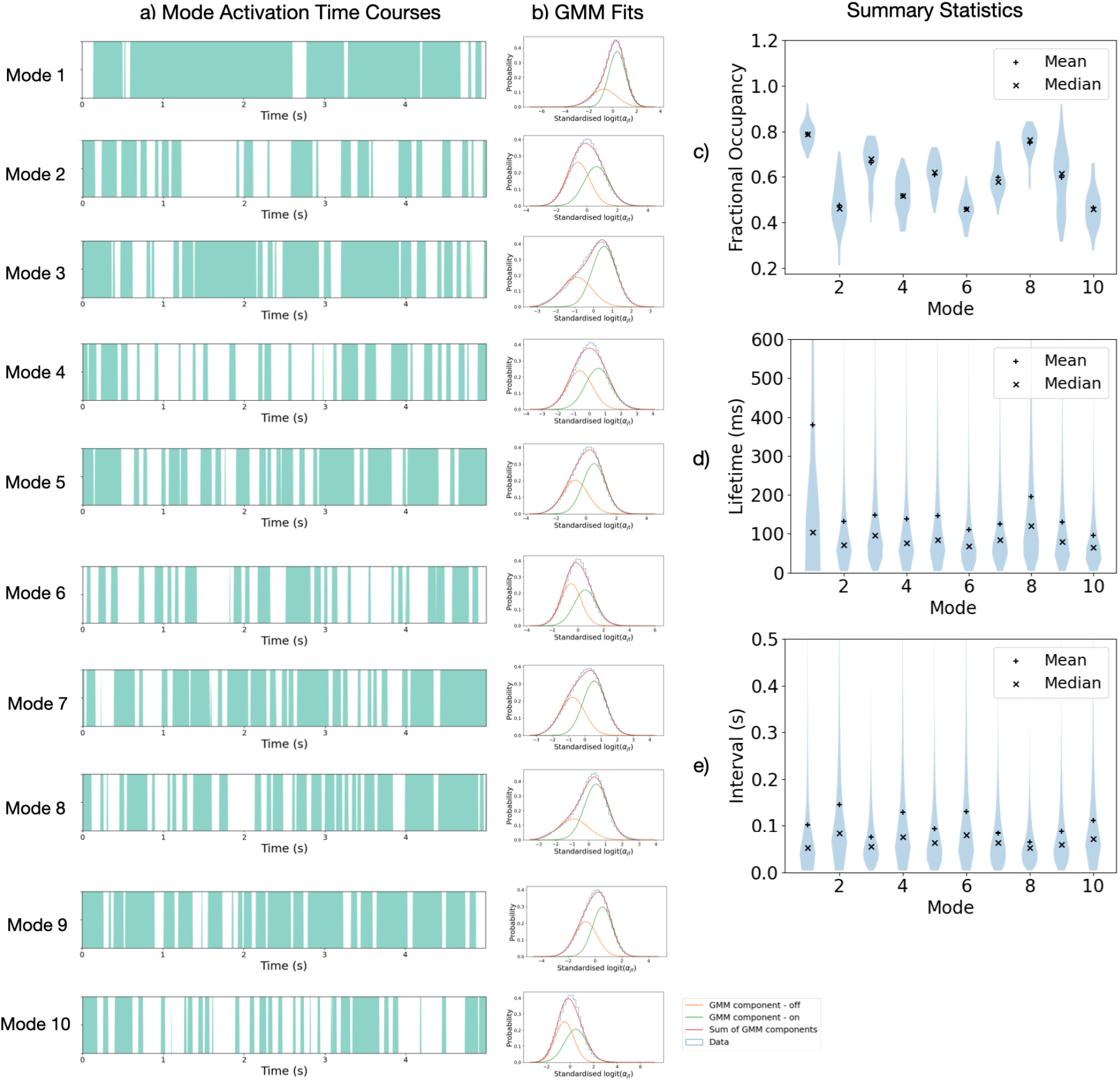
DyNeMo reveals short-lived mode activations with lifetimes of 100-150ms. a) Mode activation time courses. Turquoise regions show when a mode is “active”. Only the first 5 seconds of each mode activation time course for the first subject is shown. b) GMM fits used to identify mode “activations”. Distribution over activations and subjects of c) mode activation lifetimes and d) intervals. e) Distribution over subjects of fractional occupancies. We see mode 1 has a significantly longer mean lifetime (approximately 400 ms) compared to the other modes (approximately 100-150 ms). There is also a wide distribution of fractional occupancies across subjects.

#### DyNeMo learns long-range temporal correlations

Latent temporal correlations in MEG data can be seen by examining the inferred mixing coefficients, which are shown in Figure 8. A process is considered to possess long-range temporal correlations if its autocorrelation function decays sufficiently slowly (usually measured relative to an exponential decay) [70, 71]. The autocorrelation function and PSD form a Fourier transform pair, therefore, we can examine the presence of long-range temporal correlations by looking at the PSD. Figure 10b (top left) shows the PSD of the inferred mixing coefficients. The PSDs are rapidly decaying with a 1/*f*-like spectrum. This indicates the autocorrelation function must have a slow decay, suggesting the presence of long-range temporal correlations. As in Section 3.1, this does not mean that DyNeMo’s generative model has necessarily learnt long-range dependencies, as the presence of long-range temporal correlations could be a result of purely data-driven information. We can examine if the generative model in DyNeMo was able to learn these long-range temporal correlations by sampling a mixing coefficient time course from the model RNN. Figure 10a shows a sampled mixing coefficient time course. The PSD of the mixing coefficient time course sampled from the model RNN, Figure 10b (bottom left), shows the same 1/*f*-like spectrum as the inferred mixing coefficient time course, demonstrating it was able to learn long-range temporal correlations in the data. This is in contrast to an HMM, where the PSD of the inferred state time course, Figure 10b (top right), shows long-range temporal correlations, but the PSD of a sampled state time course, Figure 10b (bottom right), does not. It is also worth noting that the inferred long-range temporal correlations for the HMM are also less strong than for DyNeMo. This implies that the DyNeMo inferred long-range temporal correlations are not purely data driven, but also come from knowledge about long-range temporal correlations captured by DyNeMo through gathering information across the whole dataset. Note, although the HMM was not able to learn long-range temporal correlations, it was still able to infer them. This is because the inference depends on both the model and the data. Despite the limited memory in the HMM, there is sufficient information coming from the data to infer long-range temporal correlations in the states.

**Figure 10:**
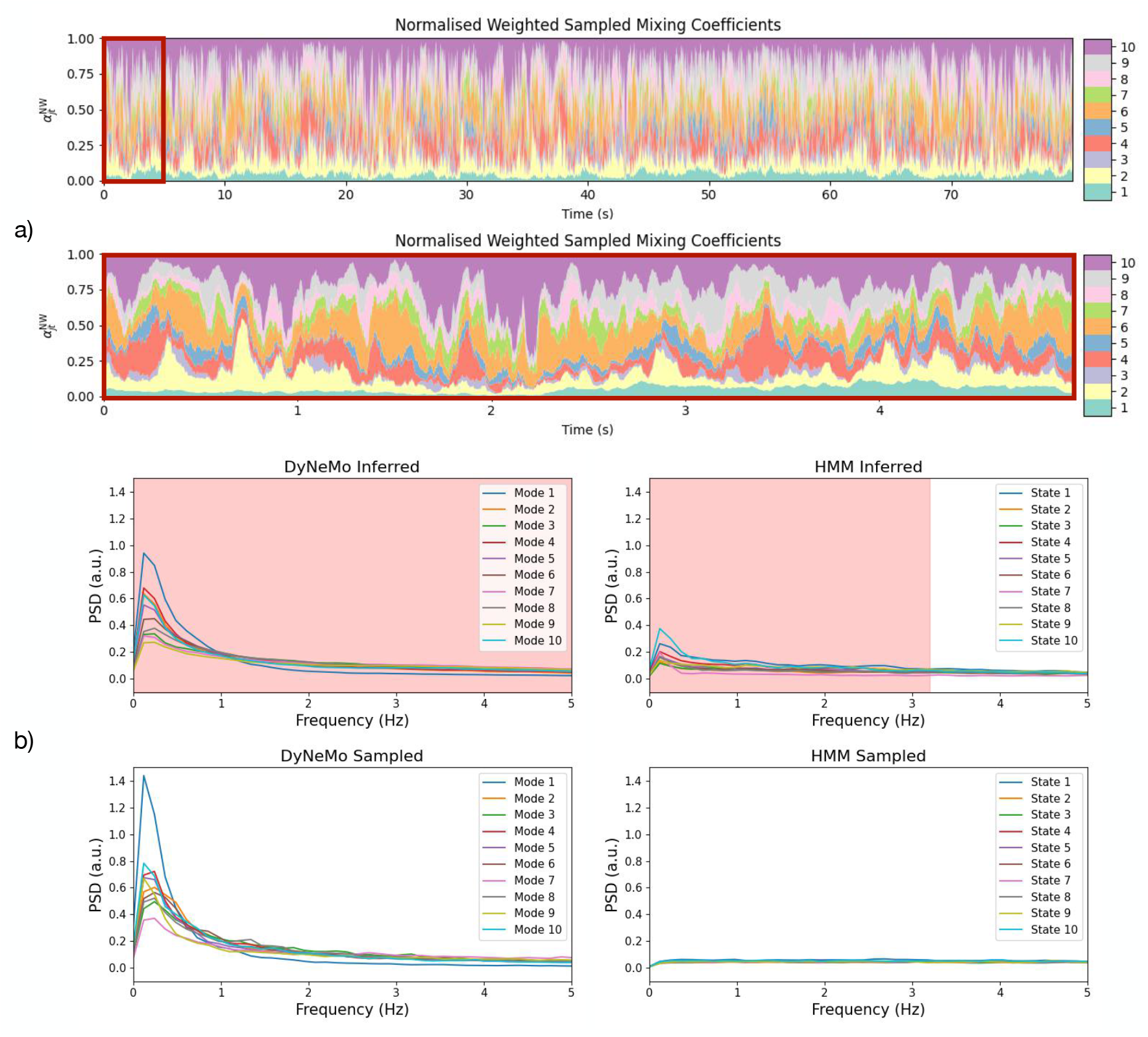
DyNeMo learns long-range temporal correlations in resting-state MEG data. a) Normalised weighted mixing coefficients sampled from the DyNeMo model RNN trained on resting-state MEG data. b) PSD of the sampled and inferred normalised weighted mixing coefficients from DyNeMo and sampled and inferred state time courses from an HMM. The red dashed line in b) shows statistically significant frequencies (p-value < 0.05) when comparing the inferred time courses with a sample from the HMM using a paired t-test. The mixing coefficient time course sampled from the DyNeMo model RNN resembles the inferred mixing coefficient time course and shows a similar PSD. Contrastingly, the sampled state time course from an HMM does not have the same temporal correlations as the inferred state time course, which is demonstrated by the flat PSD for the sample. Each mixing coefficient time course was standardised (z-transformed) across the time dimension before calculating the PSD. The fractional occupancy in a 200 ms window was used to calculate the PSD of the HMM state time courses, see [14].

#### Large-scale resting-state networks can be formed from a linear mixture of modes

The mixture model in DyNeMo allows it to construct large-scale patterns of covariance using a combination of modes with localised activity. This can be seen by comparing the modes inferred by DyNeMo with states that reveal large-scale networks inferred by an HMM. An HMM was trained on the same resting-state dataset. Power maps, FC maps and PSDs of the HMM states are shown in Figure S7. Two important networks identified by the HMM are the anterior and posterior default mode networks (states 1 and 2). The power map for DyNeMo mode 10 (see Figure 6) resembles the anterior state, however, there is no single mode that resembles the posterior state. Figure 11a shows the correlation of HMM state time courses with DyNeMo mode mixing coefficient time courses. We can see the modes that are correlated most with a state time course have activity in similar locations. Focusing on the default mode network states, DyNeMo mode 4 is the most correlated the posterior state and mode 10 is most correlated with the anterior state. In [62], it was shown that the default mode networks states have a high power in the alpha band for the posterior state and in the delta/theta band for the anterior state. The PSDs of the modes 4 and 10 also show this, providing further evidence that these modes are an alternative perspective on these states. The contribution of each mode to the default mode network HMM states is shown in Figure 11b. This shows the ratio of the total power in a mode relative to the total power in an HMM state. We can see that the power in the default mode network states is explained by many modes, i.e. DyNeMo has found a representation of these states that combines many modes. This is also true for the other HMM states. Figure 11 shows the fraction of power explained by a certain number of modes for each HMM state. The fraction of power explained increases monotonically with number of modes with no one particular mode explaining a large fraction of power. The mode description provided by DyNeMo appears to be fundamentally different to the HMM, no segments of time where one mode dominates are found. Instead, it is a representation where multiple modes co-exist and dynamics are captured by changes in the relative activation of each mode.

**Figure 11:**
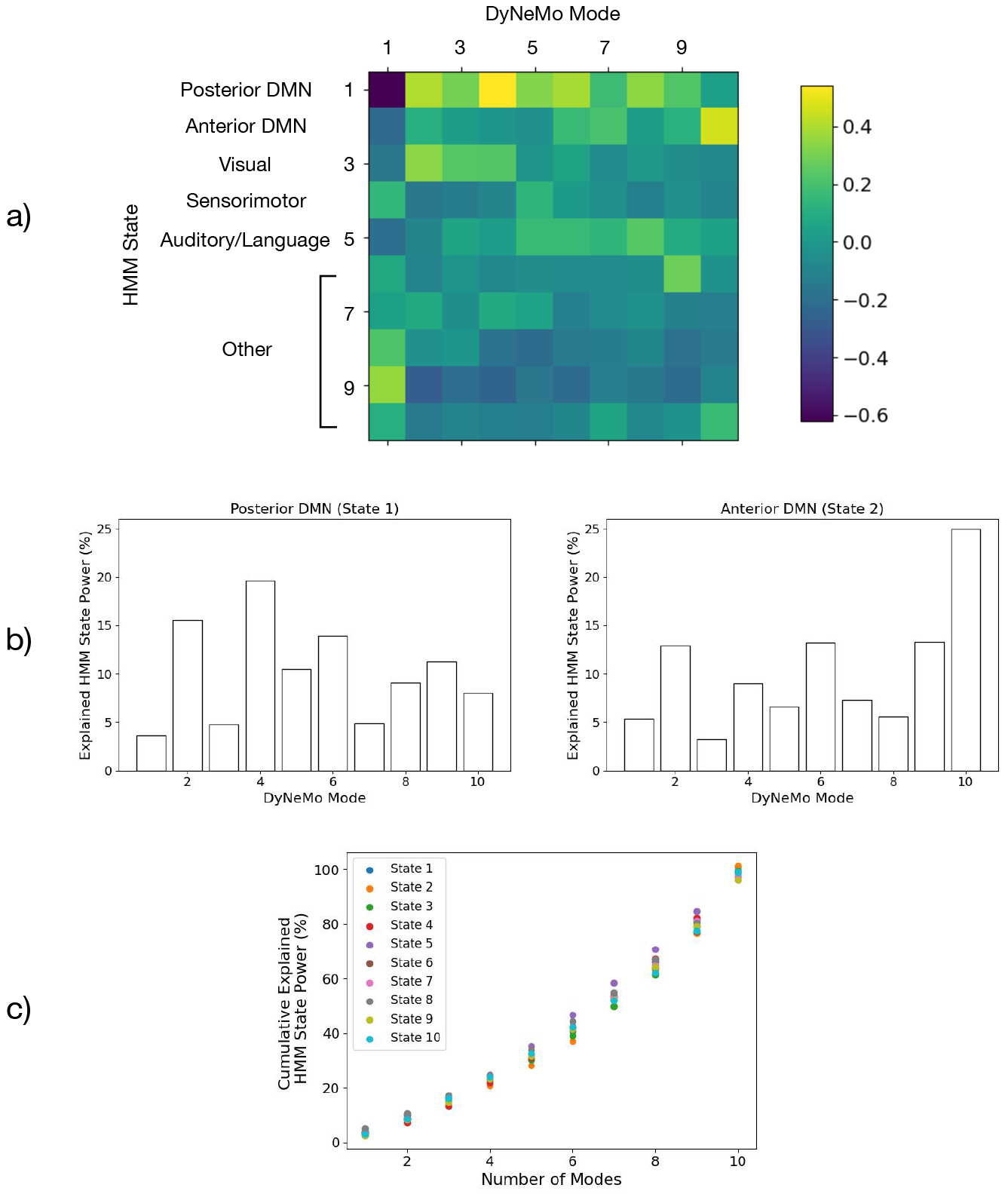
HMM states can be represented as a linear mixture of modes. a) Correlation of HMM state time courses with DyNeMo mode mixing coefficient time courses. The dynamics of multiple mode time courses correlate with each HMM state time course. In particular, many modes co-activate with the posterior default mode network (DMN) state. All elements are significant with a *p*-value < 0.05. b) Percentage of HMM state power explained by each DyNeMo mode for the posterior and anterior DMN. This was calculated as 〈*α_jt_*〉 Tr(*D_j_*)/Tr(*H_i_*), where *D_j_*(*H_i_*) is the DyNeMo (HMM) covariance for mode *j* (state *i*) and 〈*α_jt_*〉 is the time average mixing coefficient for mode *j* when state *i* is active. This shows all modes contribute to some extent to the power in these HMM states. c) The cumulative explained power for each HMM state. The modes were re-ordered in terms of increasing contribution before calculating the cumulative sum. Error bars are too small to be seen.

### 3.4 Task MEG Data

#### Resting-state networks are recruited in task

The power maps, FC maps and PSDs of 10 modes inferred by DyNeMo trained from scratch on the task MEG dataset described in Section 2.3.3 are shown in Figure 12. Very similar functional networks are found in task and resting-state MEG data (see Section 3.3). The main difference between the resting-state and task power maps is that the sensorimotor network has split into two asymmetric modes. This could be due to the more frequent activation of this area in the task dataset, which incentivises the model to infer modes that best describe power at this location.

**Figure 12:**
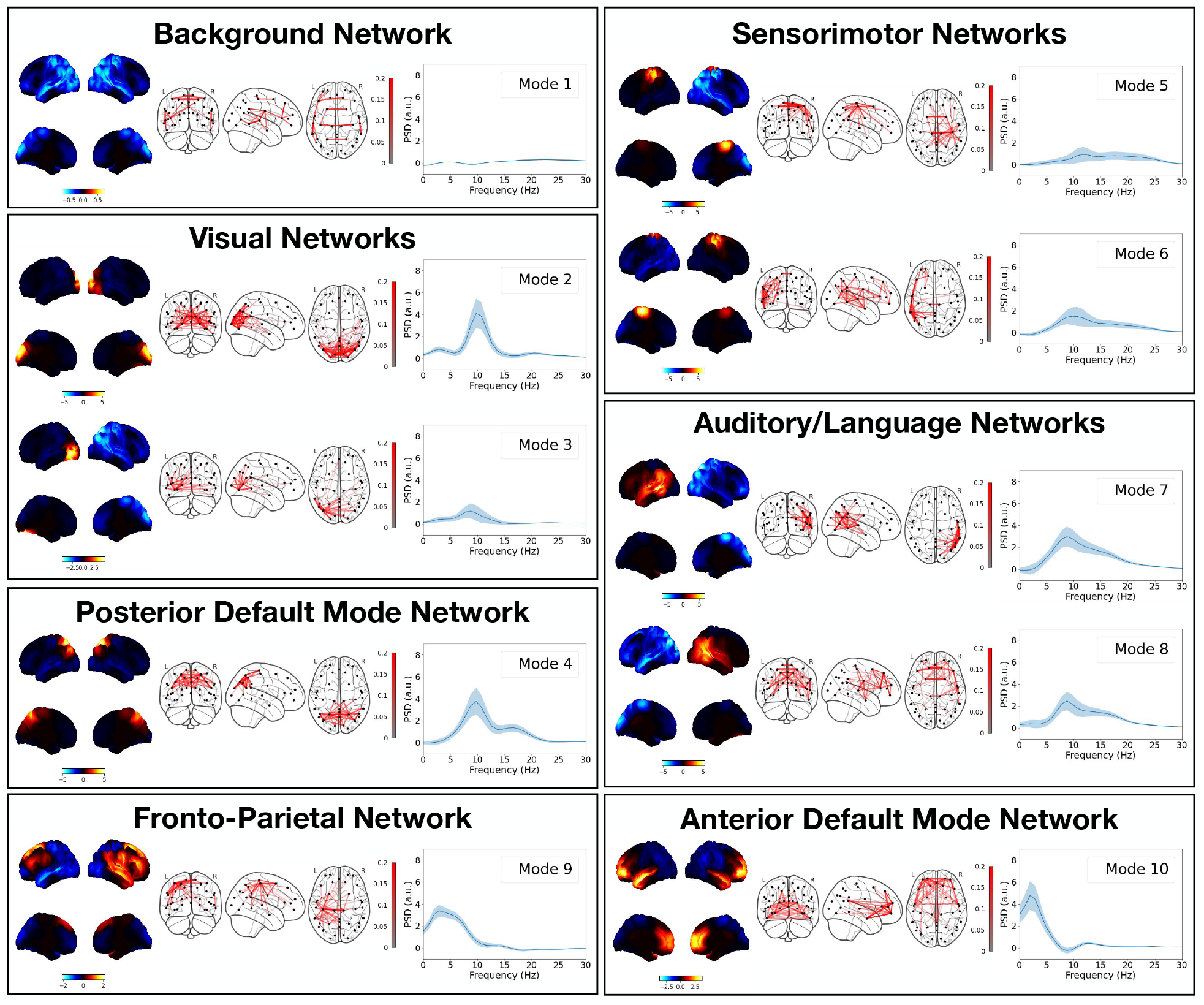
Resting-state networks are recruited in task. Ten modes were inferred using task MEG data from 51 subjects. Very similar functional networks are inferred as the resting-state data fit shown in Figure 6. Modes are grouped in terms of their functional role. Each box shows the power map (left), FC map (middle) and PSD relative to the mean averaged over regions of interest (right) for each group. The top two views on the brain in the power map plots are lateral surfaces and the bottom two are medial surfaces. The shaded area in the PSD plots shows the standard error on the mean.

#### Modes show an evoked response to task

When the inferred mixing coefficient time courses are epoched around task events, an evoked response is seen. With the window around the presentation of the visual stimulus (Figure 13a, left), DyNeMo shows a strong activation in mode 2 which corresponds to activity in the visual cortex. It also shows smaller peaks in modes 4 (posterior default mode network) and 8 (auditory/language) followed by another larger peak in mode 9 (fronto-parietal network). These represent neural activity moving from the visual cortex to a broader posterior activation and finally to an anterior activation. With the window around the abduction event (Figure 13a, right), DyNeMo shows a strong peak in mode 5, which corresponds to activity in the motor cortex. This is accompanied by a broader suppression of mode 4 which represents the posterior default mode network. The presence of task-related activations in the mixing coefficient time courses when DyNeMo is unaware of the task structure of the data demonstrates its ability to learn modes that are descriptive of underlying brain activity.

**Figure 13:**
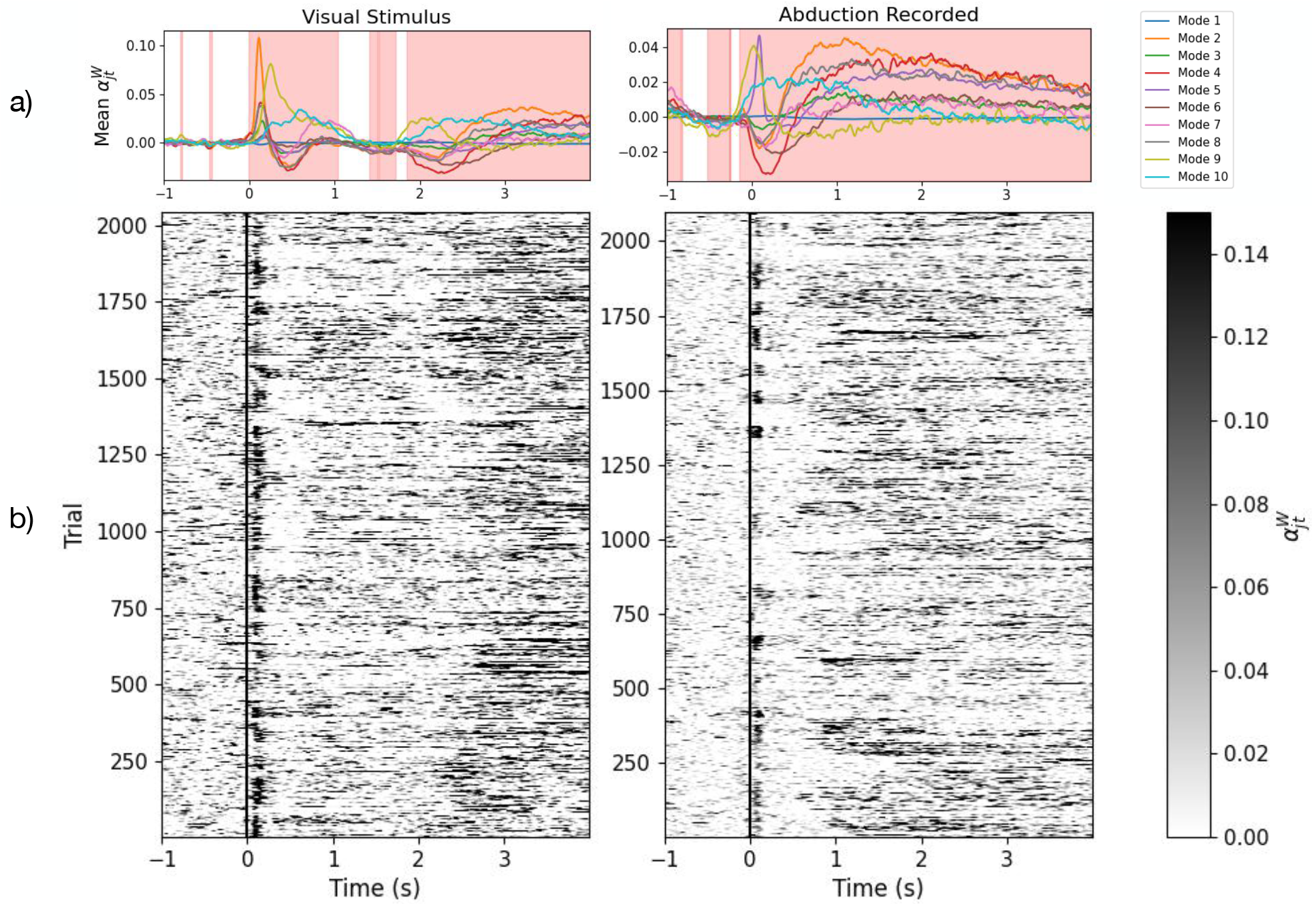
A consistent task-dependent response to the visuomotor task is seen for a number of modes. a) Trial-averaged mode timecourses weighted by the trace of their mode covariances epoched around the visual (left) and abduction (right) task. The red background shows significant time points (*p*-value < 0.05) calculated using a sign-flip permutation t-test with the family-wise error rate being controlled by using the maximum statistic. b) Individual trial responses (mode mixing coefficients weighted by the trace of their covariance) for mode 2 (visual, left) and mode 5 (sensorimotor, right). The visual stimulus/abduction task occurs at Time = 0 s.

When considering the individual trials, rather than the average response across trials, we see that the visual mode is consistently activated when the visual stimulus is presented (Figure 13b, left) and the sensorimotor mode is consistently activated when the abduction occurs (Figure 13b, right), which suggests the evoked response is not just an aggregated effect. An HMM trained on the same dataset also shows trial-wise activation (Figure S10), although the binary nature of its state activations means that the contribution of a given state can be either wiped out by another state or falsely activated by reduced activity elsewhere. DyNeMo avoids this by allowing a mixture of states to be active at a given time.

#### DyNeMo is a more accurate model of dynamic spectral properties compared to an HMM

Epoching the spectrogram of the source reconstructed data we can see the evoked response to task as a function of frequency (Figure 14). For the visual task (Figure 14a, left), immediately after the stimulus we can see a sharp increase in power around 5 Hz followed by a reduction in power around 10 Hz and above. This is repeated again around 2 s into the epoch, which is when the visual stimulus is removed. For the abduction task (Figure 14a, right), immediately after the task we also see a sharp increase in power at 5Hz followed by a reduction in power at 10 Hz and above. However, this is followed by an increase in power at 10 Hz and above, commonly known as a post-movement beta rebound [72, 73]. We can reconstruct a model estimate for the spectrogram of the data from a DyNeMo (HMM) fit by multiplying the inferred mode (state) time course by the estimate of the mode (state) PSD. Model estimate spectrograms are shown for DyNeMo and the HMM in Figures 14b and 14c respectively, along with their reconstruction errors (i.e. the residual, *ϵ*_*t*_(*f*), in Eq. (39)). The absolute value of the reconstruction error averaged over frequency for DyNeMo and the HMM is shown in Figure 14d. Both DyNeMo and the HMM are able to model dynamics in spectral content of the data, however, DyNeMo shows a modest improvement in the time-averaged reconstruction error of 5.0% (4.0%) for the visual (abduction) task compared to 5.2% (4.7%) for the HMM. A paired t-test shows the difference between the DyNeMo and HMM reconstruction error is significant with a p-value < 0.01.

**Figure 14:**
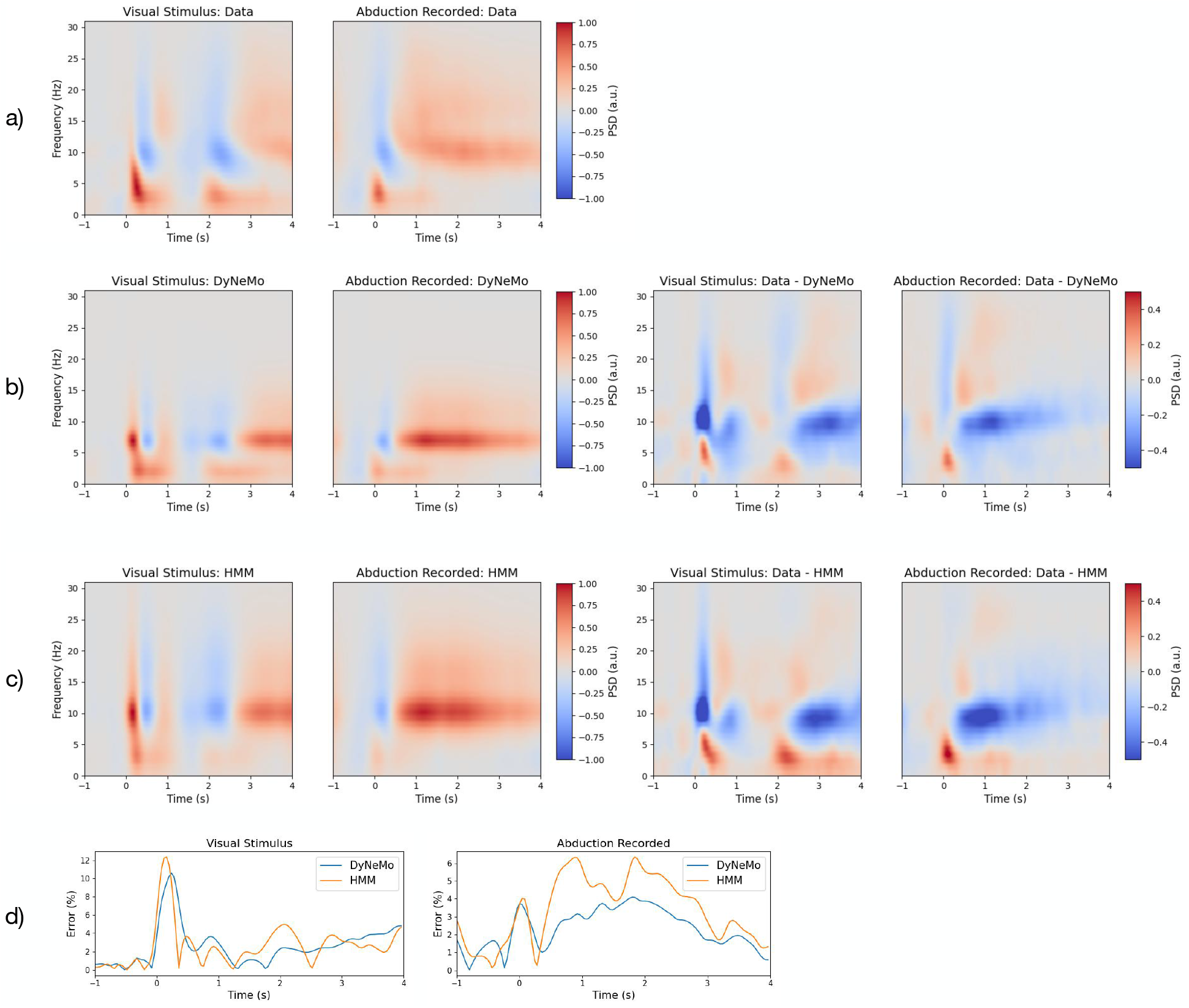
DyNeMo is a more accurate model of spectral properties compared to an HMM. a) Spectrogram of the source reconstructed data epoched around the visual and abduction task. The spectrogram was baseline corrected by subtracting the mean for the duration before the task (for each frequency separately). b) The DyNeMo model reconstruction of the spectrogram epoched around the visual and abduction task (left) and the difference from the spectrogram of the source reconstructed data (right). c) The HMM reconstruction of the spectrogram epoched around the visual and abduction task (left) and the difference from the spectrogram of the source reconstructed data (right). The spectrogram of the data and reconstruction from both models have been normalised to the range −1 to 1. The average spectrogram across all channels is shown. d) Absolute value of the reconstruction error for DyNeMo and the HMM averaged across frequencies for the visual (left) and abduction task (right). The reconstruction error is expressed as a percentage of power at each time point calculated by averaging the spectrograms in (a) over frequency. DyNeMo shows a smaller error in reconstructing the data spectrogram compared to the HMM, indicating it is a more accurate model of spectral properties.

## 4 Discussion

We have shown that MEG data can be described using multiple modes of spatiotemporal patterns that form large-scale brain networks (Figures 6 and 12). Recently, other models that provide a mode description of neuroimaging data have been proposed. Ponce-Alvarez et al. and Tewarie et al. used non-negative tensor factorisation to identify dynamic overlapping spatial patterns of connectivity [74, 36]. Núñez et al. used community detection on a time series of FC matrices to identify repeated patterns of connectivity [75]. Atasoy et al. propose ‘connectome harmonics’, where an eigendecomposition of the Laplacian of a structural connectivity matrix is calculated, which results in a set of harmonic modes that represent spatial patterns of connectivity [76]. Atasoy et al. showed that these modes predict resting-state networks [76]. Glomb et al. and Rué-Queralt et al. used the modes as a basis set to obtain a spatiotemporal description of EEG data, which revealed fast dynamics [77, 78]. Although, these technique provide a dynamic description of the data using a set of overlapping spatial modes, they all lack a generative model. Furthermore, connectome harmonics are determined from the structural connectivity matrix. In DyNeMo, a mode description of the FC is learnt directly from the data (see Section 2).

The modes inferred by DyNeMo have distinct spectral properties and correspond to plausible FC systems, such as visual, sensorimotor, auditory or other higher-order cognitive activity. These modes are more localised and can be more lateralised than the spatial patterns attributed with HMM states. Previous analysis of resting-state MEG data using an HMM [62] was able to identify large-scale transient networks which exist on time scales between 50 and 100 ms. We find DyNeMo infers transient networks at similar time scales of 100-150 ms (Figure 9). The implies the fast dynamics inferred by an HMM are not due to the assumption of mutually exclusive states.

An HMM trained on the resting-state MEG dataset used in this work suggested the default mode network was split into an anterior and posterior component [62]. In DyNeMo, the default mode network is further split up into many modes that combine to represent this network (Figure 11b). The modes that represent the default mode network show power in the same regions and frequency bands as the HMM states, supporting the fact that the modes represent an alternative perspective on the data.

Training DyNeMo on task MEG data, we find similar functional networks as inferred with resting-state data (Figure 12). This finding is supported in literature for other neuroimaging modalities, where the same networks are found in resting-state and task fMRI data [4]. The similarity in the functional networks could also be due to the fact that the majority of the subjects in the task dataset are also present in the resting-state dataset.

In an unsupervised fashion, DyNeMo was able to infer modes associated with the task. This is seen as an evoked response in the mixing coefficients of a mode to a task (Figure 14). This demonstrates that the modes inferred by DyNeMo meaningfully represent brain activity. The modes also reflect the expected time-frequency response to visual and motor tasks, which builds confidence in the description provided by DyNeMo. We find DyNeMo provides a more accurate model compared to an HMM of time-varying spectral features in the training data (Figure 14). However, both DyNeMo and the HMM show errors in modelling high-frequency spectral content in the task MEG dataset. We believe this arises from the PCA step in the data preparation, which retains components that explain large amounts of variance. In this data, lower frequencies have larger amplitudes and are able to explain more variance than high frequencies with smaller amplitudes, leading to high-frequency spectral content being filtered out. Avoiding the loss of this information could be investigated in future work with spectral pre-whitening techniques.

The smaller reconstruction error for the spectrogram of task MEG data from DyNeMo is due to the linear mixture affording the model a greater flexibility to precisely model dynamics. The fact that the reconstruction error is only slightly reduced compared to the HMM suggests that despite the constraint of mutual exclusivity the HMM was still able to provide a good description of dynamics.

### 4.1 Methodological Advancements

We believe that DyNeMo improves upon alternative unsupervised techniques in four key ways: the use of amortised inference; the use of the reparameterisation trick; the ability to model data as a linear mixture of modes (opposed to mutually exclusive states) and the ability to model long-range temporal dependencies in the data.

The amortised inference framework used in DyNeMo (described in Section 2) contains a fixed number of trainable parameters (inference RNN weights and biases). This means DyNeMo is readily trainable on datasets of varying size. Usually, the number of trainable parameters in the inference network is significantly smaller than the size of a dataset, making this approach very efficient when scaling to bigger datasets. As the availability of larger datasets grows, so does the need for models that can utilise them. Here, we believe deep learning techniques will play an important role, where with more data, models with a deep architecture begin to outperform shallower ones. Although, in this work we have studied a relatively small dataset (51-55 subjects) using a shallow model (one RNN layer), DyNeMo is readily scalable in terms of model complexity to include multiple RNN layers and more hidden units. In combination with bigger datasets this can reveal new insights into brain data. For example, previous modelling of a large resting-state fMRI dataset (Human Connectome Project, [79]) using an HMM revealed a link between FC dynamics and heritable and psychological traits [80]. The training time for DyNeMo and the computational expense of the analysis presented in this work is comparable to the HMM training time and analysis performed with the HMM-MAR toolbox^4^ presented in [62]. We believe due to the use of amortised inference, DyNeMo will be a more efficient option for larger datasets compared to the HMM-MAR toolbox.

Provided we are able to apply the reparameterisation trick to sample from the variational posterior distribution, we are able to infer the parameters for any generative model. This facilitates the use of more sophisticated and non-linear observation models and opens up a range of future modelling opportunities. This includes the use of an autoregressive model capable of learning temporal correlations in the observed data; the hierarchical modelling of inter-subject variability and the inclusion of dynamics at multiple time scales, similar to the approach used in [45].

A key modelling advancement afforded by DyNeMo is the ability to model data as a timevarying linear sum of modes. The extent to which modes mix is controlled by a free parameter referred to as the *temperature, τ*, which appears in the softmax transformation of the logits (see Equation (36) in SI 9.2). Low temperatures lead to mutually exclusive modes whereas high temperatures lead to a soft mixture of modes. In this work, we allow the temperature to be a trainable parameter. By doing this, the output of the softmax transformation is able to be tuned during training to find the appropriate level of mixing to best describe the data. Such a scheme can be interpreted as form of entropy regularisation [81, 82].

The inclusion of a model RNN in DyNeMo allows it to generate data with long-range temporal dependencies (Figures 4 and 10). This is because the future value of a hidden logit is determined by a long sequence of previous values, not just the most recent value. There is significant evidence for long-range temporal correlations in M/EEG data [83, 70, 84] and an association between altered long-range temporal correlations and disease [85, 86]. Models that are capable of learning long-range temporal correlations are advantageous in multiple ways: they can be more predictive of task or disease than models with a shorter memory; they can prevent overfitting to noise in the training data through regularisation and finally they can be used to synthesise data with realistic long-range neural dynamics.

In addition to the modelling and inference advancements discussed above, we also proposed a new method for calculating spectral properties for data described using a set of modes (see Section 2.4). With an HMM, methods such as a multitaper [12] can be used to provide high-resolution estimates of PSDs and coherences for each state. This approach relies on the state time course identifying segments of the training data where only one state is active. This approach is no longer feasible with a description of the data as a set of co-existing modes. In this paper, we propose fitting a linear regression model to a cross spectrogram calculated using the data. This method relies on different time points having different ratios of mixing between the modes. Provided this is the case, this method produces high-resolution estimates of the PSD and coherence of each mode (Figures 6, 12 and 14).

### 4.2 Drawbacks

As with most modern machine learning models, DyNeMo contains a large number of hyperparameters that need to be specified before the model can be trained. These are discussed in SI 9.2. An important hyperparameter that affects the interpretation of inferences from the model is the number of modes, *J*. We discuss the impact of varying the number of modes in SI 9.5. In short, as the number of modes is increased, the spatial activity of each mode becomes more localised and the variability of the inferred spatial patterns increases. The variational free energy is an approximation to the model evidence [47] so can be used to compare models with a different number of modes. However, Figure S4 shows the variational free energy decreases monotonically up to 30 modes. This implies more modes provide a better model for the data. As we increase the number of modes we lose the low-dimensional interpretable description of the data. Because of this trade-off we specify the number of modes by hand rather than using the variational free energy. Additionally, we ensure any conclusions that are based on studies using DyNeMo are not sensitive to the number of modes chosen. We tune other hyperparameters by seeking the set of parameters that minimise the value of the loss function.

In addition to a large number of hyperparameters, we find the model is sensitive to the initialisation of trainable parameters. This includes the internal weights and biases of RNN layers and the learnable free parameters for the mode means and covariances. The initialisations used in this work are listed in SI 9.2. We found the initialisation of the mode covariances to be particularly important. We overcome the issue of sensitivity to the initialisation of trainable parameters by training the model from scratch with different initialisations and only retaining the model with the lowest loss.

### 4.3 Outlook and Future Applications

The model presented here has many possible future applications. For example, it could be used to provide a dynamic and interpretable latent description, as done in this work, for other datasets. Alternatively, it could be used to facilitate future studies, examples of which are described below.

A common method to study the brain is the use of temporally unconstrained multivariate pattern analysis (*decoding*) to predict task, disease or behavioural traits [87]. The latent representation inferred by DyNeMo (unsupervised) provides a low-dimensional form of the training data, which is ideal for such analyses. This can overcome overfitting issues that are commonly encountered in decoding studies that use the raw data directly. Alternatively, the model architecture could be easily modified to form a semi-supervised learning problem where the loss function used has a joint objective to learn a low-dimensional representation that is useful for decoding as well as reconstructing the training data.

A useful feature of DyNeMo is the possibility of *transfer learning*, i.e. the ability to transfer information learnt from one dataset to another. This could be exercised by simply training DyNeMo on one dataset from scratch, before fine tuning the model on another dataset, which would facilitate the transfer of information through all the trainable parameters of the model, such as RNN weights, mode means/covariances, etc. Large resting-state datasets are commonplace in neuroimaging. A problem encountered in studies of small datasets (e.g. comprising of diseased cohorts) is the lack of statistical power for drawing meaningful conclusions [88]. Leveraging information gained from larger resting-state datasets could improve the predictions made on smaller datasets. For example, it has been shown resting-state data is predictive of task response [89, 90]. We believe DyNeMo offers the possibility of transferring information acquired from resting-state datasets with thousands of individuals to the individual subject level.

The generative model proposed here explictly models the covariance of the training data as a dynamic quantity. In this paper, we trained on prepared (time-delay embedded/PCA) source reconstructed data. However, the model could be trained on unprepared sensor-level data to estimate the sensor covariance as a function of time. Such a model could be utilised in the field of M/EEG source reconstruction. Algorithms for source reconstruction often assume the sensor-level covariance is static, which is rarely the case [91]. Using a dynamic estimate of the covariance, we can construct time-vaying reconstruction weights for source reconstruction [40], which can improve source localisation.

Finally, whilst we focused on parcellated source reconstructed MEG data in this paper, DyNeMo could of course be applied to data from other neuroimaging modalities such as fMRI, sensor level MEG data and other electrophysiological techniques (EEG, ECOG, etc.).

## 5 Conclusions

We have proposed a new generative model and accompanying inference framework for neuroimaging data that is readily scalable to large datasets. Our application of DyNeMo to MEG data reveals fast transient networks that are spectrally distinct, in broad agreement with existing studies. We believe DyNeMo can be used to help us better understand the brain by providing an accurate model for brain data that explicitly models its dynamic nature using a linear mixture of modes. The modest improvement in modelling dynamic spectral properties compared to an HMM shows the assumption of mutual exclusivity does not necessarily impact the HMM’s ability to model the data effectively. Nevertheless, DyNeMo is a novel and complementary tool that is useful for studying neuroimaging data.

## 6 Acknowledgments

We would like to thank Matt Brookes and his team at the University of Nottingham for providing us with the data analysed here. Data were collected in the context of the Medical Research Council (MRC)-funded MEG UK partnership. We would also like to thank Yan-Ping Zhang-Schaerer for her feedback in developing DyNeMo.

This research was supported by the National Institute for Health Research (NIHR) Oxford Health Biomedical Research Centre. The Wellcome Centre for Integrative Neuroimaging is supported by core funding from the Wellcome Trust (203139/Z/16/Z). C.G. is supported by the Wellcome Trust (215573/Z/19/Z). E.R. is supported by an Engineering and Physical Sciences Research Council (EPSRC) and MRC grant (EP/L016044/1) and F. Hoffmann-La Roche. R.T. is supported by the EPSRC and MRC (EP/L016052/1). C.H. is supported by the Wellcome Trust (215573/Z/19/Z). A.Q. is supported the MRC (RG94383/RG89702) and by the NIHR Oxford Health Biomedical Research Centre. U.P. is supported by an MRC Mental Health Data Pathfinder award (MC/PC/17215). J.vA. is supported by EPSRC (EP/N509711/1) and Google DeepMind. P.N. is supported by EPSRC Industrial Cooperative Awards in Science & Technology (18000077) and GlaxoSmithKline. Y.G. holds a Turing AI Fellowship (Phase 1) at the Alan Turing Institute, which is supported by the EPSRC (V030302/1). M.W. is supported by NIHR Oxford Health Biomedical Research Centre, the Wellcome Trust (106183/Z/14/Z and 215573/Z/19/Z), and the New Therapeutics in Alzheimer’s Diseases (NTAD) study supported by the MRC and the Dementia Platform UK.

## 7 Ethics Statement

All participants gave written informed consent and ethical approval was granted by the University of Nottingham Medical School Research Ethics Committee.

## 9 Supplementary Information (SI)

### 9.2 Derivation of the Loss Function

In variational Bayesian inference we infer a parameter by minimising the variational free energy,

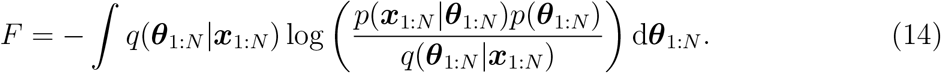

where *q*(***θ***_1:*N*_|***x***_1:*N*_) is the posterior, *p*(***θ***_1:*N*_) is the prior and *p*(***x***_1:*N*_|***θ***_1:*N*_) is the likelihood, ***θ***_*t*_ is the logit at each time point, ***x***_*t*_ is the observed data at each time point and *t* = 1,…,*N* denotes the time index. We can separate the logarithm into two terms,

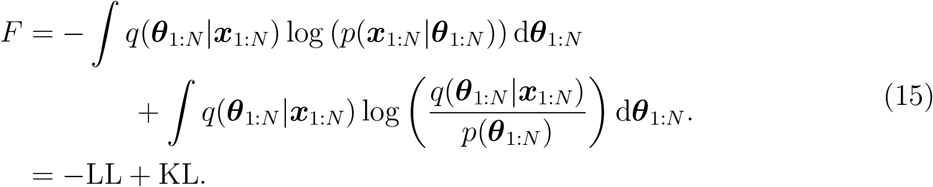

LL is referred to as the *log-likelihood term* and KL is referred to as the *KL divergence term*. Considering the log-likelihood term,

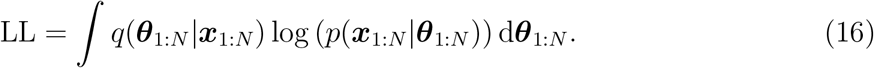

We use the mean field approximation for the posterior,

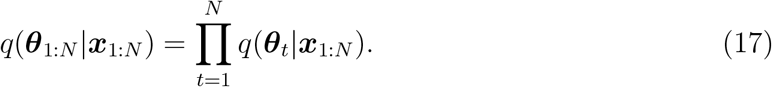

Each factor *q*(***θ***_*t*_|***x***_1:*N*_) is a multivariate normal distribution parameterised by a mean vector ***m***_*θ*_*t*__ (***x***_1:*N*_) and diagonal covariance matrix 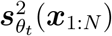, i.e.

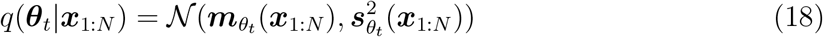

We also assume the data at each time point is independent and only depends on the logit at that time point, i.e. we factorise the likelihood as

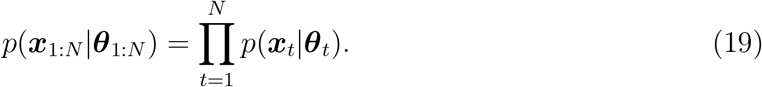

We assume a multivariate normal distribution for the data, i.e.

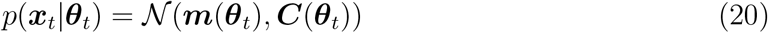

Substituting Equations (17) and (19) into Equation (16),

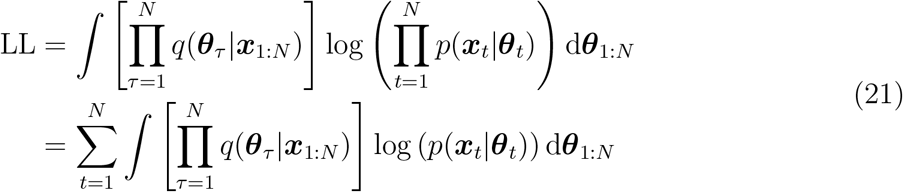

For each term in the summation we can factorise the integral as

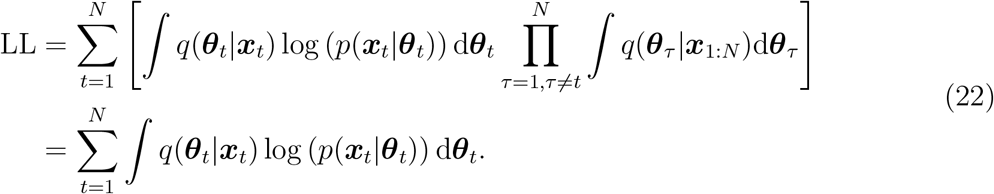

We can use a Monte Carlo estimate to calculate this as

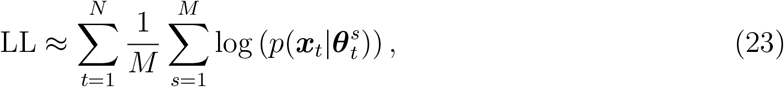

where 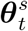 denotes the *s*^th^ sample from the posterior distribution *q*(***θ***_*t*_|***x***_1:*N*_) at time point *t*. In practice we use just one sample, i.e. *M* = 1. Therefore, the log-likelihood term is approximated by

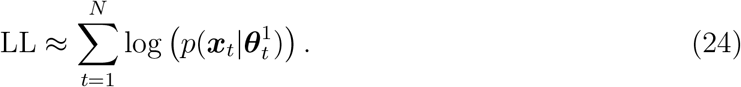

Considering the KL divergence term,

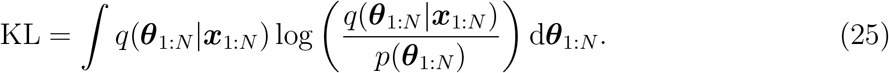

We use the mean field approximation for the posterior (Equation (17)) and factorise the prior as

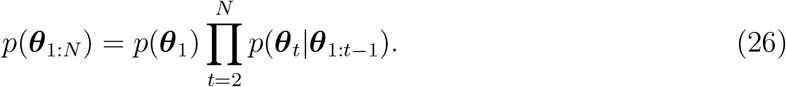

With these substitutions the KL divergence term becomes

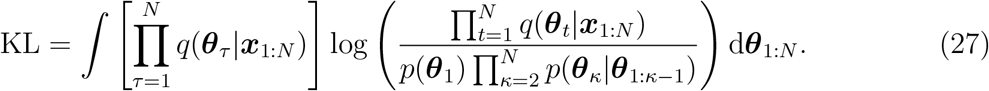

We split up the logarithm as

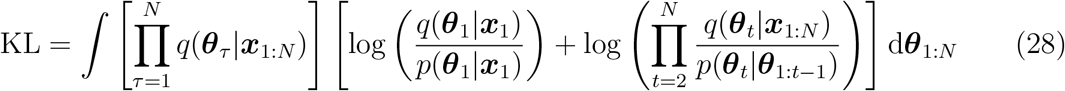

and clip the first logarithm to give

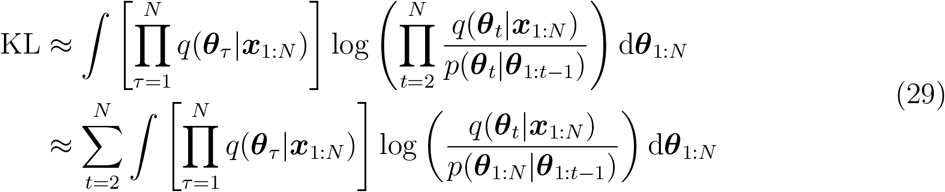

For each term in the summation, we can factorise the integral as

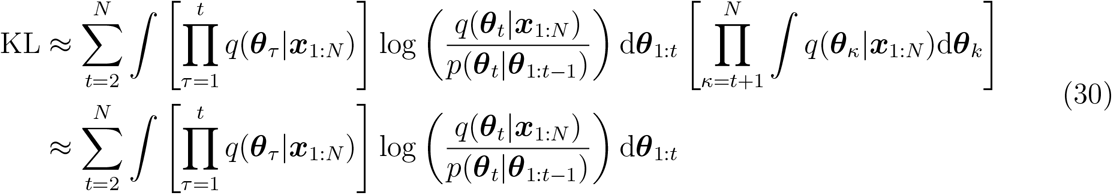

We denote the integral over d***θ***_*t*_ by

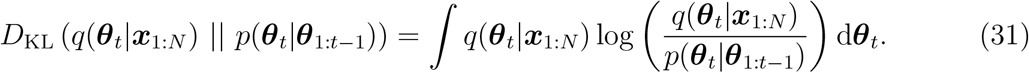

Substituting this into the KL divergence term, we get

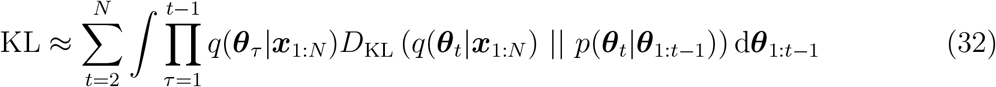

We use a Monte Carlo estimate using a single sample from each posteriors *q*(***θ***_1_|***x***_1:*N*_),…, *q*(***θ***_*t*-1_|***x***_1:*N*_). Therefore, our KL divergence term is

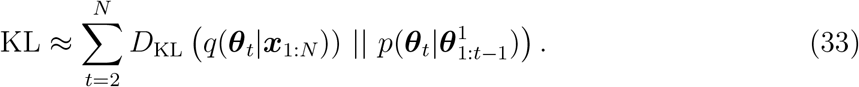

The prior 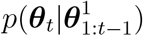 is a multivariate normal distribution parameterised by a mean vector 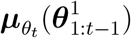 and diagonal covariance matrix 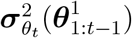, i.e.

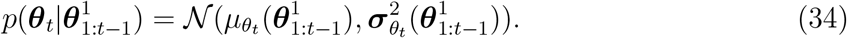

The parameters 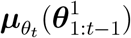 and 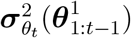 are calculated using the model RNN.

We use stochastic gradient descent to minimise a loss function. The loss function we use

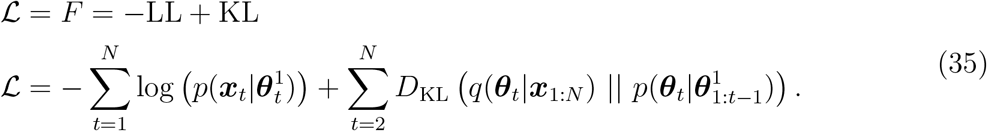

Using this loss function will minimise the variational free energy, or equivalently, it will maximise the evidence lower bound [51].

### 9.2 Training and Hyperparameters

Before training the model we prepare the source reconstructed data. We applied time-delay embedding, PCA and standardisation. Time-delay embedding and PCA are summarised in Figure S1. The procedure used to train the model and choices for hyperparameters are discussed below.

**Figure S1:**
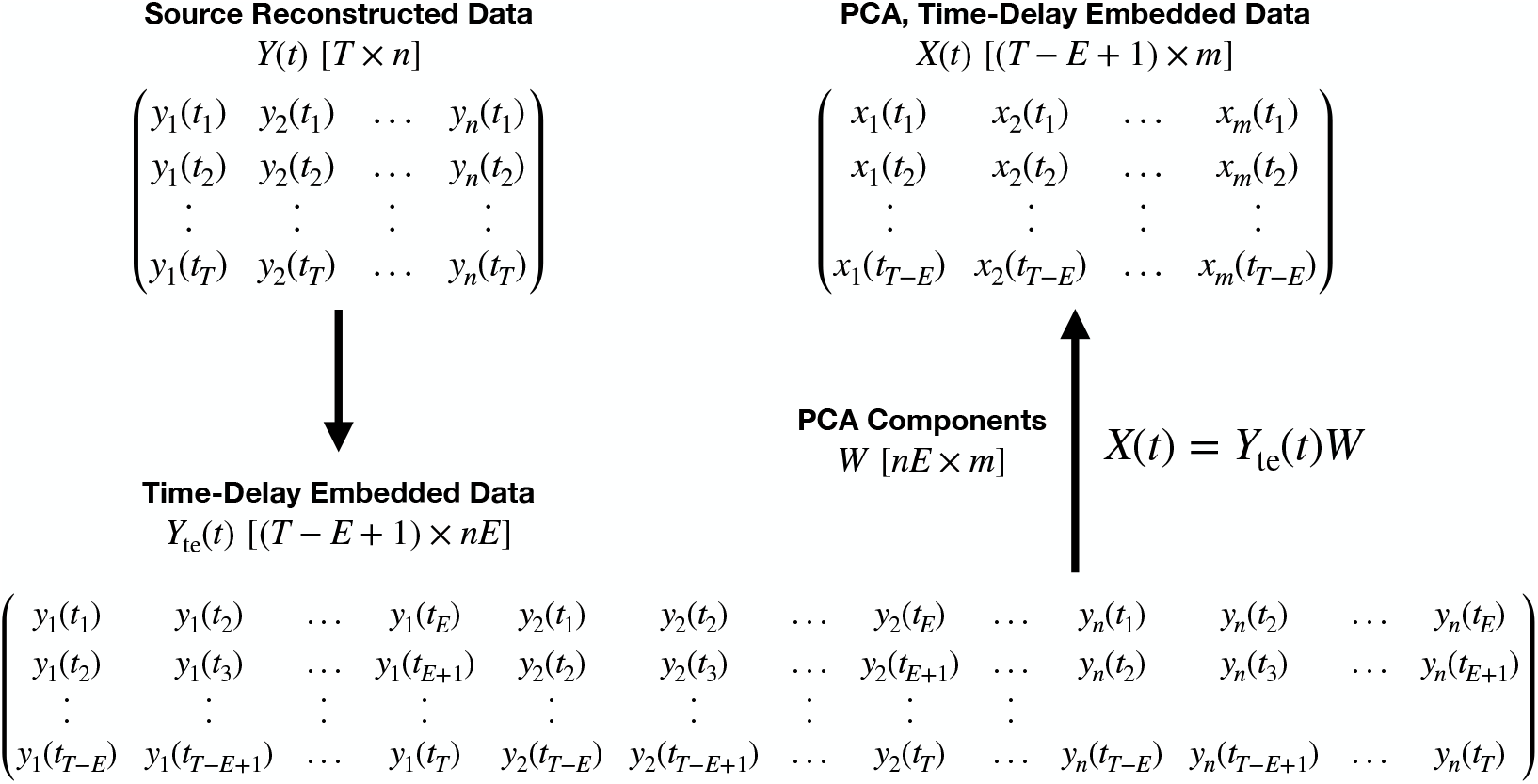
Preparation applied to the source reconstructed data before training. The use of time-delay embedding encodes spectral properties of the training data into the covariance by adding extra elements that correspond to the auto-correlation function to the matrix. PCA is performed to reduce the number of channels so that the data is not too large to fit within GPU memory. Standardisation is also applied after PCA. *T* is the number of time points, *n* is the number of parcels/regions of interest, *E* is the number of time-delay embeddings and *m* is the number of channels after PCA.

#### Logit activation function

To calculate the mixing coefficients from the logits we use a softmax function,

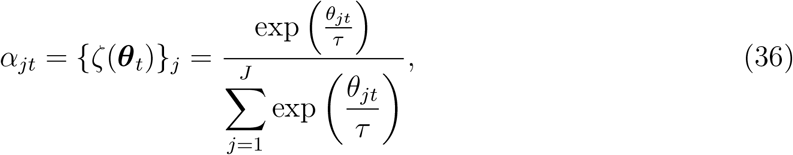

where *τ* is a hyperparameter known as the temperature, discussed further below. The use of a softmax function imposes the constraint that the *α_jt_*-values are positive and the sum of *α_jt_* over *j* is equal to one.

#### Alpha temperature

We can specify a temperature *τ* for the softmax activation function. The temperature determines the amount of mixing between modes. A high temperature corresponds to a more even mixture, whereas a low temperature leads to more mutually exclusive modes. In this work, we allow the temperature to be a learnable parameter.

#### Learning rate

In this work, we use the Adam optimiser [93] to update trainable parameters. The learning rate *η* is a key hyperparameter of the Adam optimiser, which can affect training. Using a value that is too high can lead to divergence in the loss function, alternatively, one that is too small can lead to slow convergence.

#### Sequence length

The input to the model is a sequence of data points ***x***_1:*N*_, where N is the length of the sequence. The sequence length determines the number of previous time points the model has access to. Therefore, we would like to use the longest sequence length possible. However, the sequence length is limited by the GPU memory that is available and the ability to train the model in a reasonable time frame.

#### Batch learning

Stochastic gradient descent is performed by estimating the loss for a small group of sequences, referred to as a *batch*. This loss is used to update the trainable parameters before the loss for a new batch is calculated. The number of sequences in a batch is referred to as the *batch size*. We calculate the loss for a batch by averaging the loss for each sequence in the batch. When performing batch learning it is important to shuffle the ordering of the sequences and batches. We did this by first separating the entire dataset into sequences, shuffling the order of the sequences, then grouping sequences in batches and performing one final random reordering to give the training dataset. One training loop through all of the batches is referred to as an *epoch*. The batches are not reshuffled between epochs.

#### Hidden units and number of LSTM layers

When using an LSTM we must specify a number of hidden units. We found small networks had more stable training than large networks, so only used 64 units in the model and inference LSTM. We also found stacking multiple LSTMs did not improve the model, so in this work we only use one LSTM layer.

#### Dropout

[94]. This is well-known technique used when training a neural network to mitigate overfitting. The use of dropout was found not to benefit the model so no dropout layers have been used in this work.

#### Normalisation layers

[95, 96]. Normalisation layers are often used to train deep neural networks because they help alleviate the the vanishing/exploding gradient problem [97]. In this work, we include a single Layer Normalisation [96] transformation to the output of the LSTMs in Equations (2) and (6) before the affine transformation.

#### Gradient clipping

RNNs can suffer from exploding gradients when backpropagation occurs through each time step. A strategy proposed in [98] to avoid this is gradient clipping, where we rescale the gradients so that their norm is a particular value when gradient norm would otherwise exceed this value. This strategy has been shown to improve training stability. In this work, we use gradient clipping when training on real MEG data.

#### Trainable parameters and initialisation

The trainable parameters in this model are:

- The weights and biases of the model and inference LSTM. At the start of training these are set randomly using Glorot initialisation [99].
- Layer Normalisation weights. The output of Layer Normalisation is centred around a learnable β-parameter and scaled with a γ-parameter [92, 96], these parameters were initialised using zeros and ones respectively.
- Dense layer weights and biases for the affine transformations. Glorot initialisation [99] was used for these parameters.
- The alpha temperature. This was initialised using a value of one.
- Elements of the mode means and covariances. In this work, we use zero vectors for the means and an identity matrix for the covariances. When training on MEG data we found the initialisation of the mode covariances to be important. We propose a strategy of initially training the model on the data for a randomly selected single subject to estimate its covariances. The covariances are initialised with the identity matrix when training on a single subject. This removes the subject-to-subject variability in the data. Then, the model can be trained on the full dataset initialising with the single-subject mode covariances. We found this strategy helped to avoid local optima when learning a high latent dimensionality (e.g. J > 10).

#### Multi-start training

We find the model is sensitive to the initialisation of the trainable parameters, in particular the inference and model RNNs. When training on real MEG data, we observe that the model can converge to different local optima with the same dataset. To help find the global optimum, we propose a multi-start approach, where we train the model for a small number of epochs a few times and performing the full training on the model with the lowest loss at the end of the initial training period. This procedure was used when training on real MEG data and was found to reduce the run-to-run variability.

#### KL annealing

[55] is a technique used at the start of training. An annealing factor λ is introduced into the loss function,

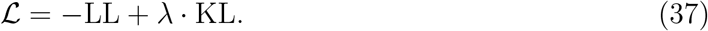

The annealing factor is a smoothly varying function of the number of training epochs. It takes a value between zero and one. The function used to calculate the annealing factor in this work is

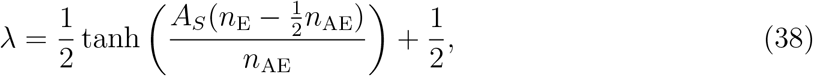

where *A_S_* is the annealing sharpness, which determines the shape of the annealing curve, *n*_E_ is the number of training epochs and *n*_AE_ is the number of annealing epochs.

At the start of training, the weights and biases of the model RNN are initialised with samples from a uniform distribution (Glorot initialisation [99]). As a consequence, the model RNN is prone to giving naive outputs/estimates for the logit time course in these early epochs. KL annealing with λ close to zero helps in this period because it prevents the model RNN from influencing the inference RNN before it has learnt a logit time course that is useful for describing the training data. As training progresses, λ tends to one and the model RNN learns the temporal dynamics in the latent representation by predicting probability distribution of the next logit ***θ***_*t*_ from the samples of previous logits ***θ***_1:*t*-1_. By doing this, it also regularises the inferred logits for the training data.

### 9.3 Post-hoc Analysis of Learnt Latent Variables

Once trained, DyNeMo provides us with a mixing coefficient time series for each mode, *α_jt_*. We use the inferred mixing coefficients with the source reconstructed data (before preparation) to perform post-hoc analysis. We describe the quantities calculated in our post-hoc analysis below.

### Summary statistics

We can summarise the mixing coefficients with statistics, which can give a high-level description of the data. We can take inspiration from the Viterbi path (referred to as the *state time course*) of an HMM^5^ [51]. We typically summarise this with state lifetimes^6^, interval times and the fractional occupancies [14]. One benefit of the mutual exclusivity assumption made by the HMM is that there are well defined time points when a state is active, making determining a lifetime and interval time straightforward. Contrastingly, DyNeMo provides a description where multiple modes are simultaneously present at each time point. To define when a mode is active we fit a two-component GMM to the mixing coefficient time series of each mode. One of the Gaussian components corresponds to time points when the mode is active whereas the other component corresponds to time points when the mode is inactive. This GMM therefore gives us a mode activation time course, which we can use to compute the usual summary statistics we would calculate with a state time course. Note, we fit a GMM separately to each mode, which enables the possibility of there being time points where multiple modes or no modes activate. The mixing coefficients have a sum to one constraint, which means their distribution is non-Gaussian. Therefore, we transform the data using the logit function^7^ and standardise before fitting the GMM to make the distribution more Gaussian. We found defining activations with a GMM led to more stable summary statistics for each mode compared to a simpler approach such as using an argmax operation. This is because with an argmax operation the mode with the largest mixing coefficient depends on the full set of modes, whereas with the GMM approach, each mode is studied in isolation.

### Mode spectral properties

Neuronal activity in the brain has oscillatory dynamics. A useful quantity for examining these oscillations is the power spectral density (PSD), which displays the power at each frequency for a given channel (i.e. region of interest). Additionally, the cross spectral density, which displays the power coupling across two channels, is of interest. We can calculate power and cross spectra for each mode directly from source reconstructed data once we have inferred the mixing coefficients.^8^ Equation (4) defines a linear mixture of mode covariances. A property of this model is that the PSD of each mode mixes with the same coefficients. We can exploit this property to estimate the spectral properties of each mode. We do this by first calculating a cross spectrogram using the dataset and fitting a linear regression model using the mixing coefficients:

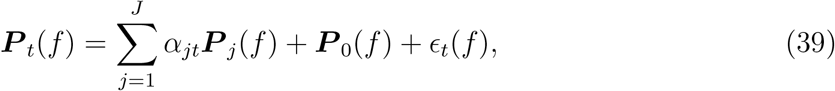

where ***P***_*t*_(*f*) is the cross spectrogram of each pair of channels, ***P***_*j*_(*f*) are regression coefficients, which are our mode cross spectra, ***P***_0_(*f*) is a mean term, *ϵ_t_*(*f*) is a residual and *f* is the frequency. We standardise (z-transform) the mixing coefficients across the time dimension before calculating the linear regression. This results in the mean term ***P***_0_(*f*) corresponding to the time-averaged PSD and the regression coefficients ***P***_*j*_ (*f*) corresponding to PSDs relative to the time-averaged PSD. We calculate the cross spectrogram with the source reconstructed data using Welch’s method [100], which involves segmenting the data into overlapping windows and performing a Fourier transform:

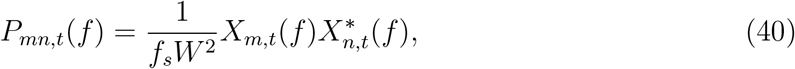

where *m* and *n* are channels, *f_s_* is the sampling frequency, *W* is the number of samples in the window, *X_m,t_*(*f*) is the Fourier transform of the data in a window centred at time *t* and * denotes the complex conjugate.

### Mode power maps

Additionally, the spatial distribution of power across the brain is of interest. We can examine the spatial power distribution of each mode separately, which can highlight the areas of the brain that are active for each mode. Using the cross spectrogram regression method (Equation (39)), we obtain a PSD for each channel for a given mode. As each channel corresponds to a region of interest, we obtain a PSD for the activity at each region of interest. The integral of a PSD is the power. Plotting the power at each region of interest as a two-dimensional heat map projected onto the surface of the brain shows the spatial distribution of power for a given mode. It is often more interesting to plot the power relative to a reference rather than the absolute value to highlight differences in the power maps of each mode. In this work, we calculate the power by integrating the mode PSDs ***P***_*j*_(*f*) without the mean term ***P***_0_(*f*). This gives us the power distribution relative to the mean power common to all modes. We also subtract the mean power across modes for each region of interest separately when displaying the power maps to help highlight relative differences between the modes. No thresholding is applied to the power maps. Additionally, the surface plotting function we use interpolates the power between parcels for visualisation.

### Mode FC maps

In addition to power maps, FC maps reveal the brain networks that are present for each mode. We use the coherence as our measure of FC, which quantifies the stability of phase difference between two regions of interest, thereby providing a measure of synchronisation or phase-locking between brain regions. We choose this measure because it gives us a direct estimate of oscillatory synchronicity, which has been proposed as a mechanism for neuronal communication [15]. To calculate the coherence we use the mode cross spectra estimated with the linear regression model in Equation (39). Estimating the coherence for a window requires us to average multiple estimates of the cross spectra within that window. We do this by dividing each window into a set of sub-windows and taking the average cross spectra for each sub-window. Using the cross spectra we calculate a coherence for each pair of channels. However, most of these connections correspond to a background level of activity that is are common to all modes. To identify the most prominent connections, we fit a two-component GMM to the distribution of coherences relative to the mean. We standardise the relative coherence values before fitting the GMM. One component of the GMM corresponds to a population of background coherence, whereas the other corresponds to prominently high relative coherence values. We plot the connection edges in the original coherence matrix that correspond to high relative coherence values. If we are unable to identify two components, we plot the top 5% of connections.

To calculate power and FC maps, we use a window length of 4 seconds, sub-window length of 0.5 seconds, window step size of 0.08 seconds, and apply a Hann windowing function to each sub-window before performing the Fourier transform to calculate the cross spectrogram. We calculate the cross spectrogram and perform the linear regression separately for each subject. We then average the subject-specific mode PSDs and coherences over subjects to obtain group-level spectra. We also apply a Hann function to sub-windows of the mixing coefficient time series to match the windowing applied to the data and average the values across a full window to calculate the *α_jt_* value to use in the regression. To calculate the spectrograms used to the study the evoked response to task (Section 3.4) we use a window length of 0.5 seconds and do not separate the window into sub-windows.

**Figure S2:**
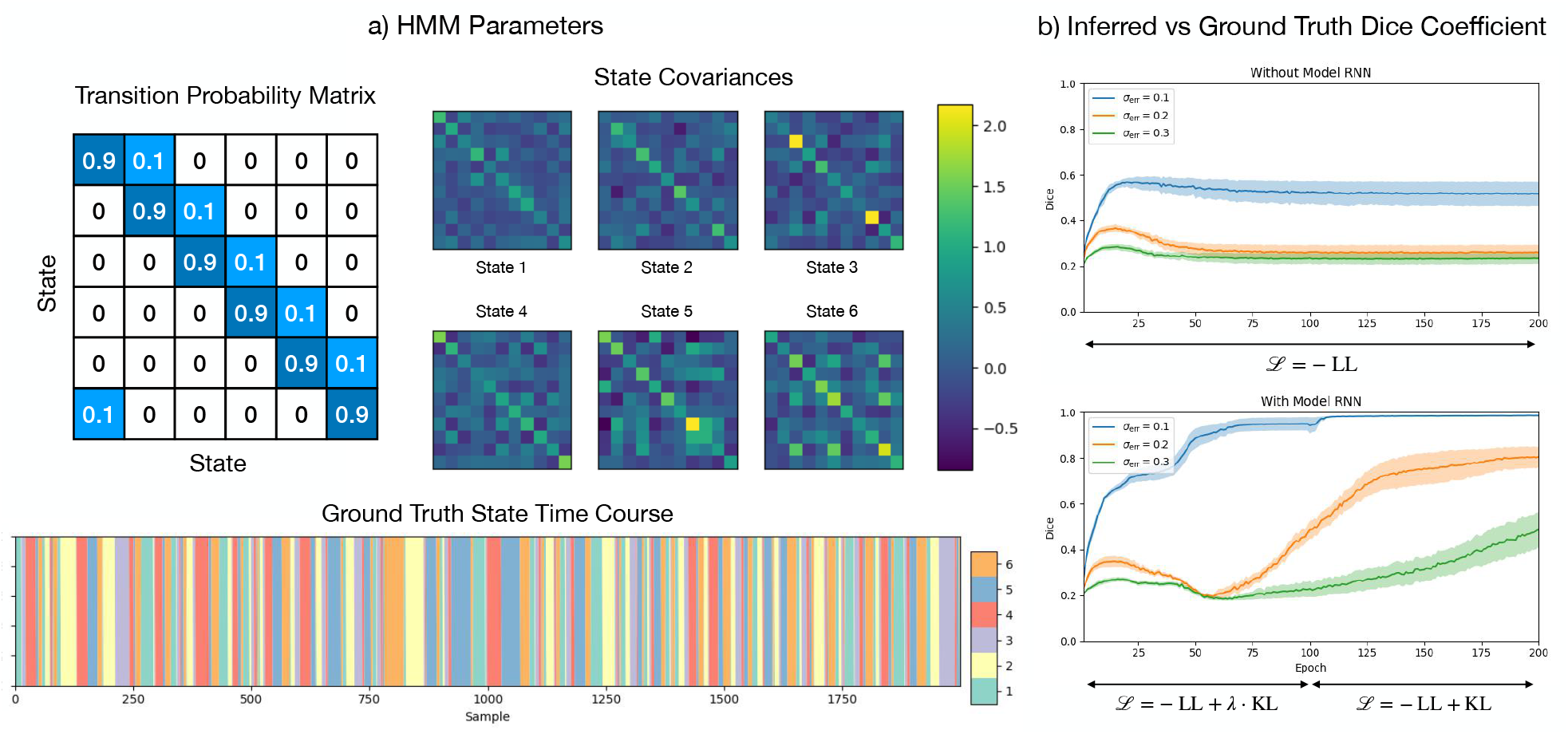
Temporal regularisation from the model RNN improves inference. a) HMM parameters used to simulate data. A 6 state, 11 channel HMM was simulated with the transition probability matrix shown. 25,600 samples were generated. Only the first 2,000 time steps of the state time course are shown. b) Dice vs epoch when minimising the negative log-likelihood loss only (top) and the variational free energy (bottom) for different added noise, *σ*_err_, which is the standard deviation of normally distributed errors added to each channel at each time point. 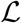 denotes the loss function used for each stage of training (see Eqs. (37) and (38)). Each model was fitted 5 times to the same dataset. The solid line is the mean and the shaded area shows the standard error on the mean.

### 9.4 Regularisation from the Model RNN

It is possible to construct a model for the training data without including the model RNN. For example, we could construct a standard variational autoencoder [48], where a normal distribution with zero mean and unit variance takes the place of the model RNN (i.e. the prior). Alternatively, point estimates for logits could be learnt and no prior would be needed. So why include a model RNN? There are two reasons, firstly learning a generative model in itself is useful, and secondly the model RNN regularises the inferred logits and can help alleviate overfitting to noise in the training dataset. The latter can be demonstrated by training DyNeMo on simulated HMM data with varying noise added. Parameters of the HMM simulation are shown in Figure S2a. We measure the inference accuracy of the model by calculating the dice coefficient of the inferred state time course and the ground truth from the simulation. Figure S2b shows the dice vs epoch during training for different levels of noise. The top figure is the achieved dice coefficient when minimising the negative log-likelihood loss only 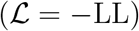. When the noise is high the model struggles to correctly infer the ground truth state time course. The bottom figure is the achieved dice when minimising the variational free energy 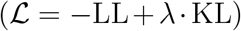. KL annealing was applied during training for the first 100 epochs. The influence of the model RNN starts to appear when λ is sufficiently large, which is after the 50th epoch roughly. Figure S2 shows the inclusion of the model RNN leads to a better inference when training on noisy data.

### 9.5 Run-to-Run Variability and Number of Modes

When training DyNeMo, our objective is to the minimise the loss function (variational free energy) by updating the trainable parameters of the model. Due to the random initialisation of the RNN weights and the use of stochastic gradient descent to update the parameters, it is possible for DyNeMo to converge to different local optima in the loss function. Neuroimaging data itself is complex and it is reasonable to expect there could be multiple models with different parameters that can lead to similar loss values. In this section we look at the variability in the final loss value for different training runs and look at the impact of varying the number of modes.

First, we look at the simulation datasets. In Bayesian inference the model evidence, which is approximated by the variational free energy [47], can be used to compare models with different hyperparameters. This allows us to select the number of modes to infer using the variational free energy. Figure S3 shows the variational free energy of DyNeMo trained on the simulation datasets described in Sections 2.3.1 and 2.3.2 as a function of number of modes. The ground truth number of modes was 3 for simulation 1 and 6 for simulation 2. We can see the variational free energy decreases until the correct number of modes is reached. It then flattens, we see adding more modes no longer shows an improvement. This indicates the variational free energy can be used to correctly select the number of modes.

Turning to the resting-state MEG dataset, we evaluate the run-to-run variability in the loss. Figure S4 shows the final training and validation loss as a function of number of modes. As the number of modes increases so does the variability in the loss. Our strategy for finding the global optimum involves training DyNeMo multiple times and selecting the model with the lowest loss. Although there’s a systematic offset in the validation loss compared to the training loss, which suggests overfitting to the 45 subject dataset, the difference remains the same as the number of modes is increased, which means overfitting does not get worse with more modes. The systematic offset is likely to be due to the fact that the validation dataset contained unseen subjects and the model not generalising to new subjects very well. Over this range, the loss decreases monotonically meaning more modes provide a better description of the data. Unlike the simulation studies, we cannot see a clear minimum in the loss function. This indicates the optimum number of modes maybe greater than 30. However, the aim of this model is to provide a low-dimensional interpretable description of the data. Therefore, rather than using the variational free energy to determine the number of modes, we preselect a low value as a hyperparameter.

**Figure S3:**
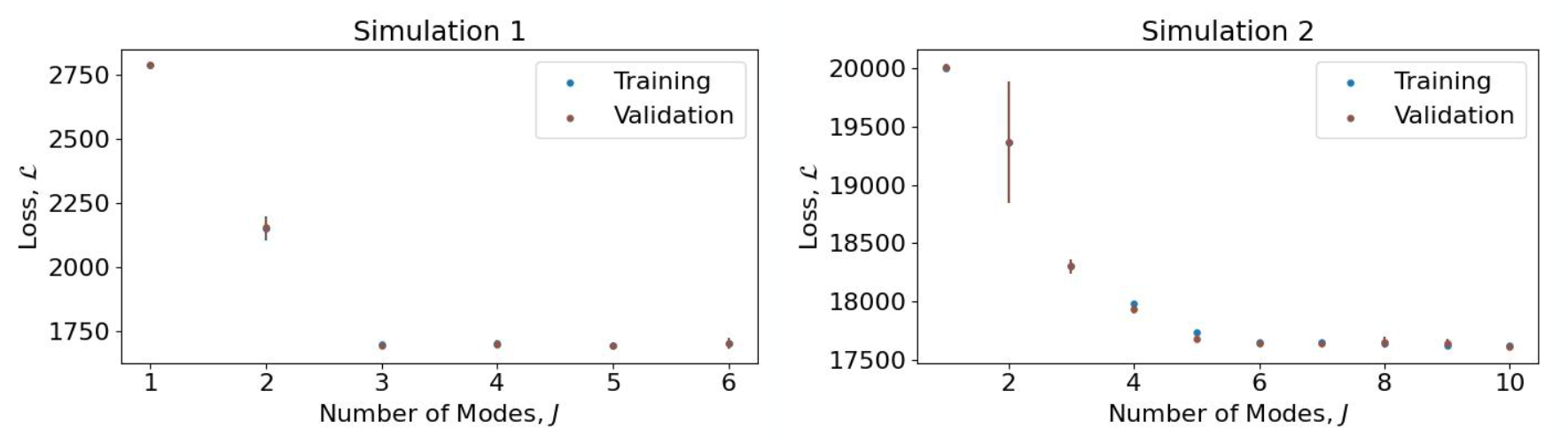
The loss (variational free energy) indicates the optimum number of modes. Simulations 1 and 2 are described in Section 3.1 and 3.2. The ground truth in these simulations was dataset generated using 3 and 6 modes respectively. The mean training (blue) and validation loss (brown) for a batch vs the number of modes. Each data point is the average of five runs. The error bar is one standard deviation. The same training dataset was used here and the results shown in Figures 4 and 5. An additional 5,120 data points were simulated for the validation dataset.

**Figure S4:**
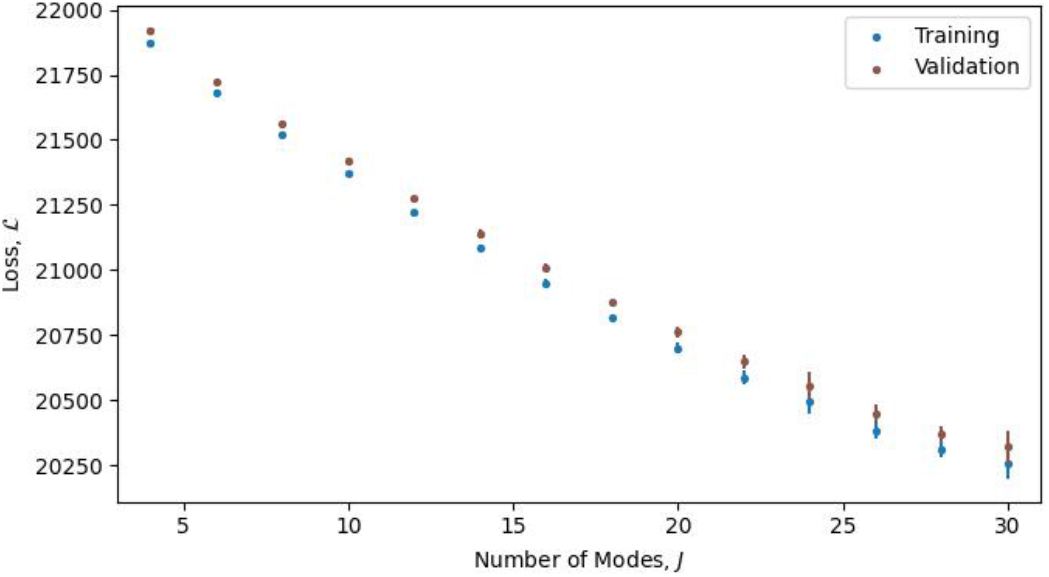
More modes provide a better description of the data. The mean training (blue) and validation loss (brown) for a batch vs the number of modes. We split the resting-state MEG dataset into a 45 subject training dataset and 10 subject validation dataset. Each data point is the average of ten runs. The error bar is one standard deviation. For 25 modes or less the error bar is too small to be seen.

Figure S5 shows the power maps for the run with the best loss when fitting 4-12 modes. As the number of modes increases the activation of each brain region becomes more localised. This is expected as increasing the number of modes allows greater precision in reconstructing the time-varying covariance ***C***_*t*_ from the mode covariances ***D***_*j*_ (see Equation (4)). When we fit 4 modes, we see power is divided amongst the four lobes: the occipital, parietal, temporal and frontal lobe. As we increase the number of modes, the number of possible ways to distribute power amongst the modes also increases. In this work, we choose to fit 10 modes to MEG data as a trade off between obtaining an interesting description of the data and minimising the variability in the power maps of each run.

**Figure S5:**
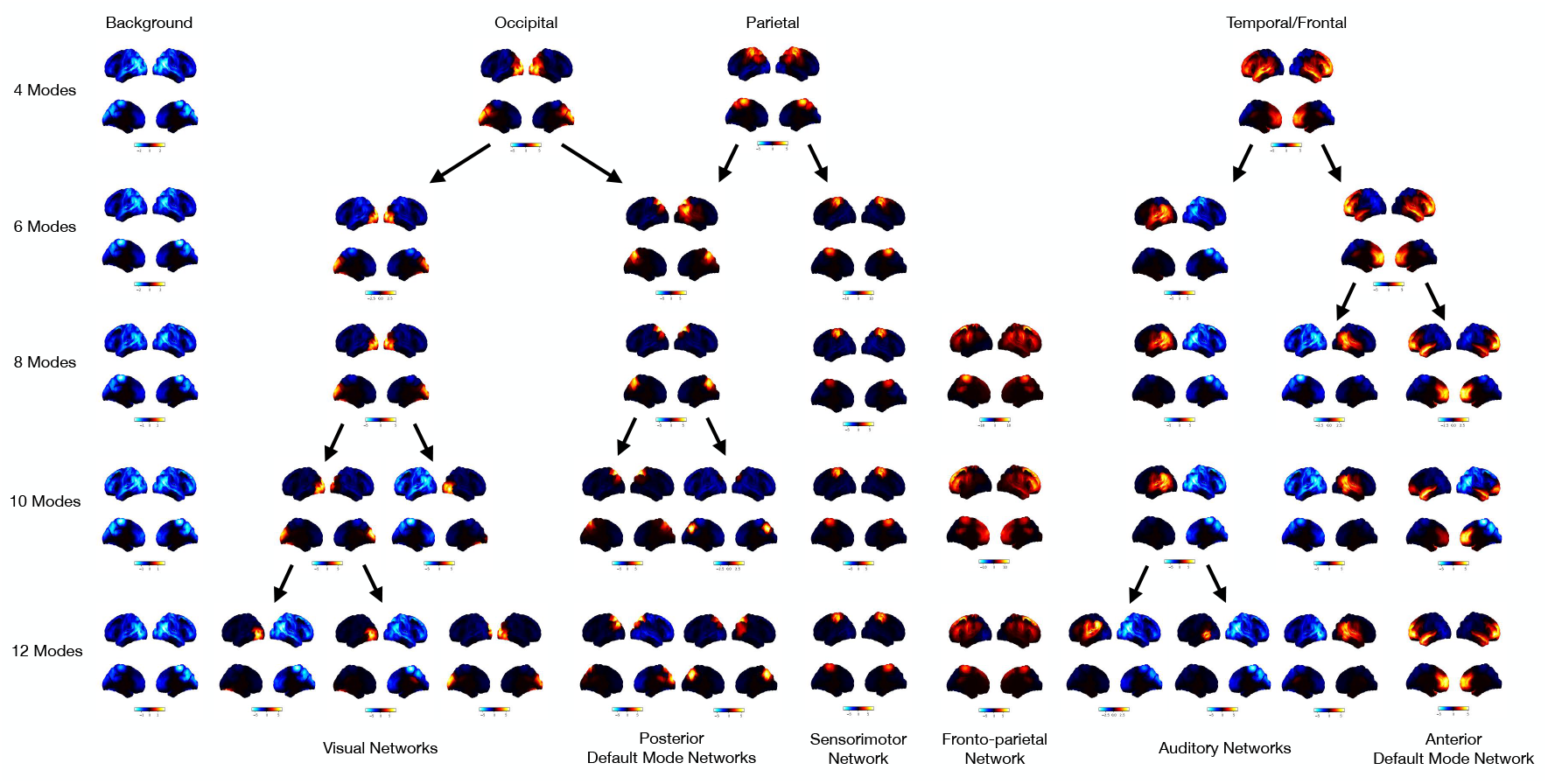
Power maps show more localised activation with an increasing number of modes. These power maps were obtained from a model trained on resting-state MEG data from 45 subjects. Arrows indicate when a power map has split as the number of modes was increased. The top two views on the brain in the power map plots are lateral surfaces and the bottom two are medial surfaces.

**Figure S6:**
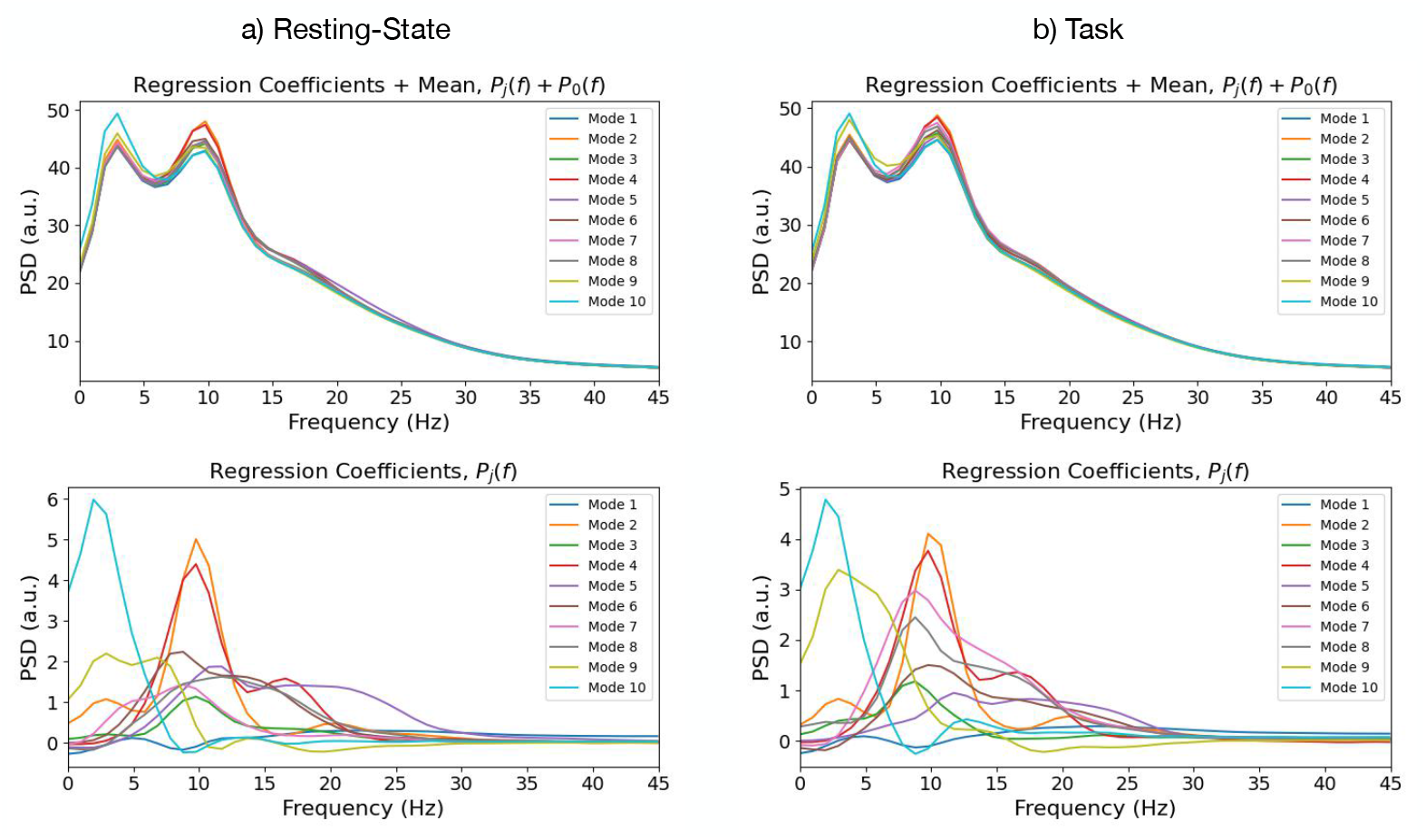
DyNeMo learns spectrally distinct modes. Mode PSDs including the activity common to all modes (top) and mode PSD relative to the mean activity common to all modes (bottom) for the resting-state (a) and task (b) MEG dataset. The mean PSD across channels is shown.

### 9.6 Mode PSDs

Figure S6 shows the PSD of each mode calculated with the regression method described in Section 2.4 for the resting-state and task MEG dataset. In Figure S6 (top) we see all modes have a PSD with a 1/f profile and exhibit a prominent 10 Hz peak. The drop off towards 0 Hz is due to filtering applied during preprocessing. In Figure S6 (bottom) we see the mode PSD relative to the activity shared across modes. We see differences in the PSD of each mode consistent with expected frequency content of activity at locations shown in the corresponding power maps.

### 9.7 HMMs Trained on the MEG Datasets

The HMM was also fitted to the MEG datasets described in Section 2.3.3 for comparison with DyNeMo. Figure S7 shows the power maps, FC maps and PSDs of the HMM states inferred on the resting-state MEG dataset and Figure S9 shows the power maps, FC maps and PSDs of the HMM states inferred on the task MEG dataset.

**Figure S7:**
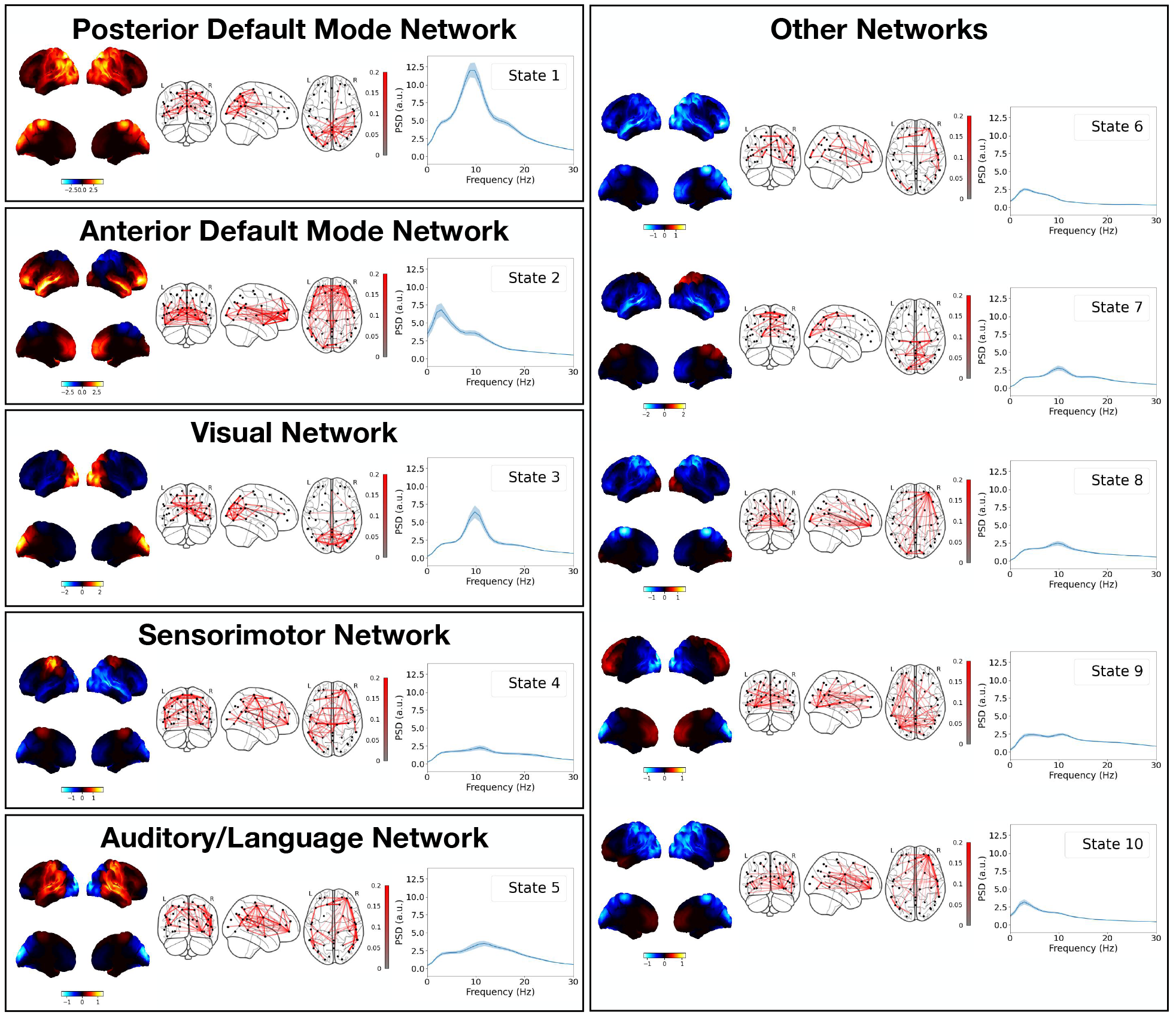
HMM states inferred on a resting-state MEG dataset consisting of 55 subjects. Each box shows the power map (left), FC map (middle) and PSD averaged over regions of interest (right) for each group. The top two views on the brain in the power map plots are lateral surfaces and the bottom two are medial surfaces. The shaded area in the PSD plots shows the standard error on the mean.

**Figure S8:**
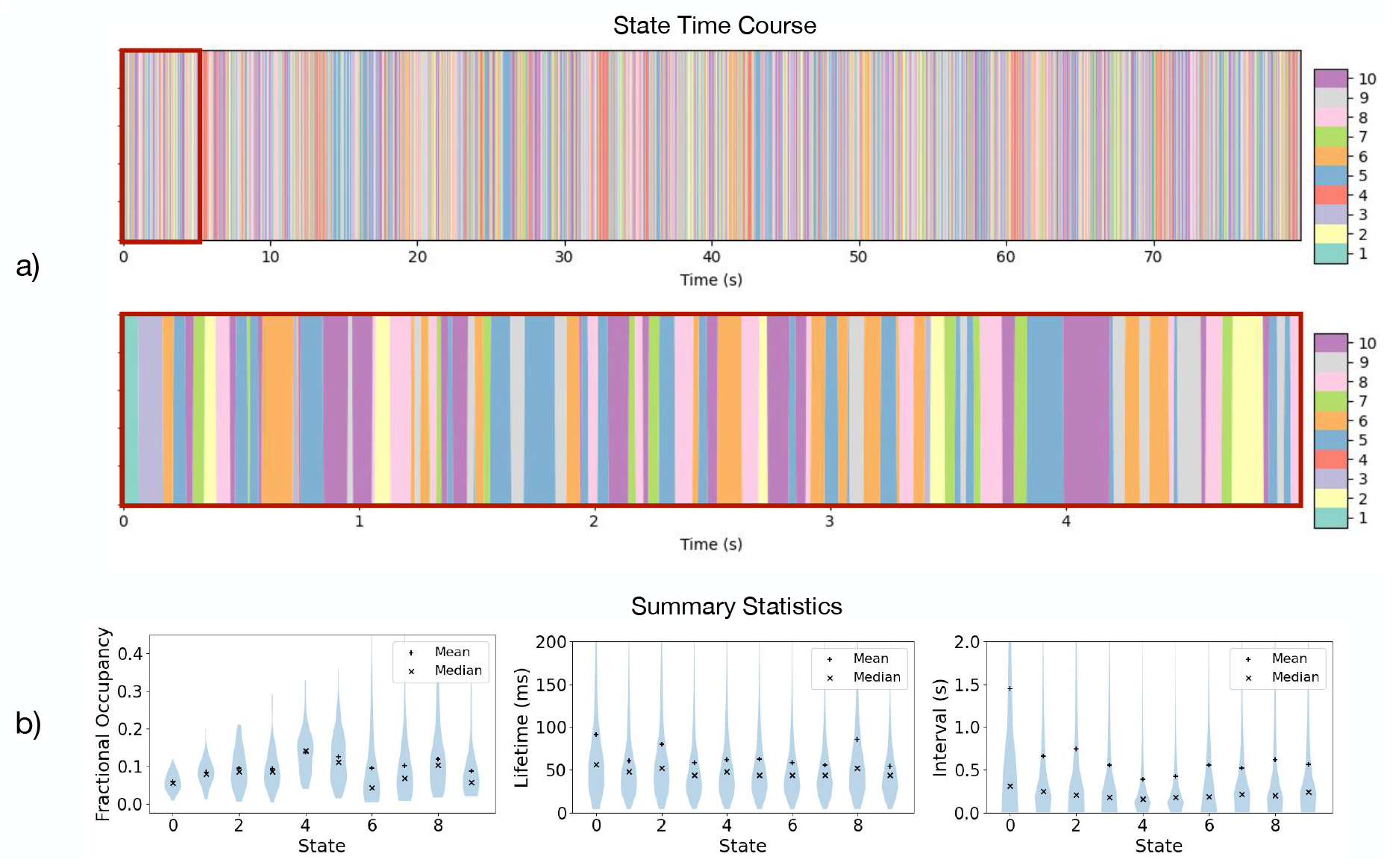
State time course and summary statistics for the HMM fit to the resting-state MEG dataset. a) Inferred state time course for the first 80 seconds of the first subject (top) and zoomed in on the first 5 seconds (bottom). b) Distribution of fractional occupancies over subjects and distribution of state lifetimes and intervals for all subjects.

**Figure S9:**
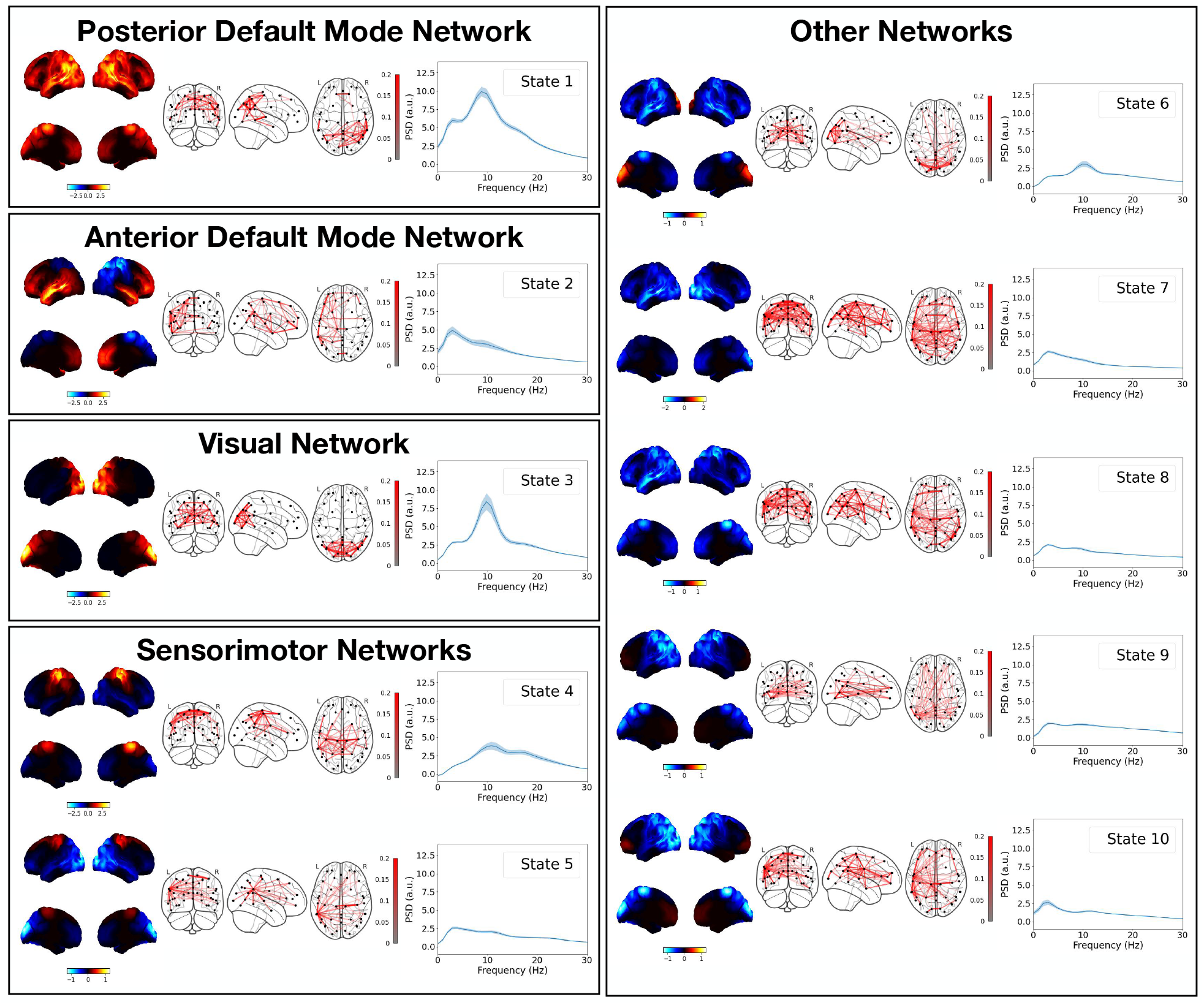
HMM states inferred on a visuomotor task MEG dataset consisting of 51 subjects. Each box shows the power map (left), FC map (middle) and PSD averaged over regions of interest (right) for each group. The top two views on the brain in the power map plots are lateral surfaces and the bottom two are medial surfaces. The shaded area in the PSD plots shows the standard error on the mean.

**Figure S10:**
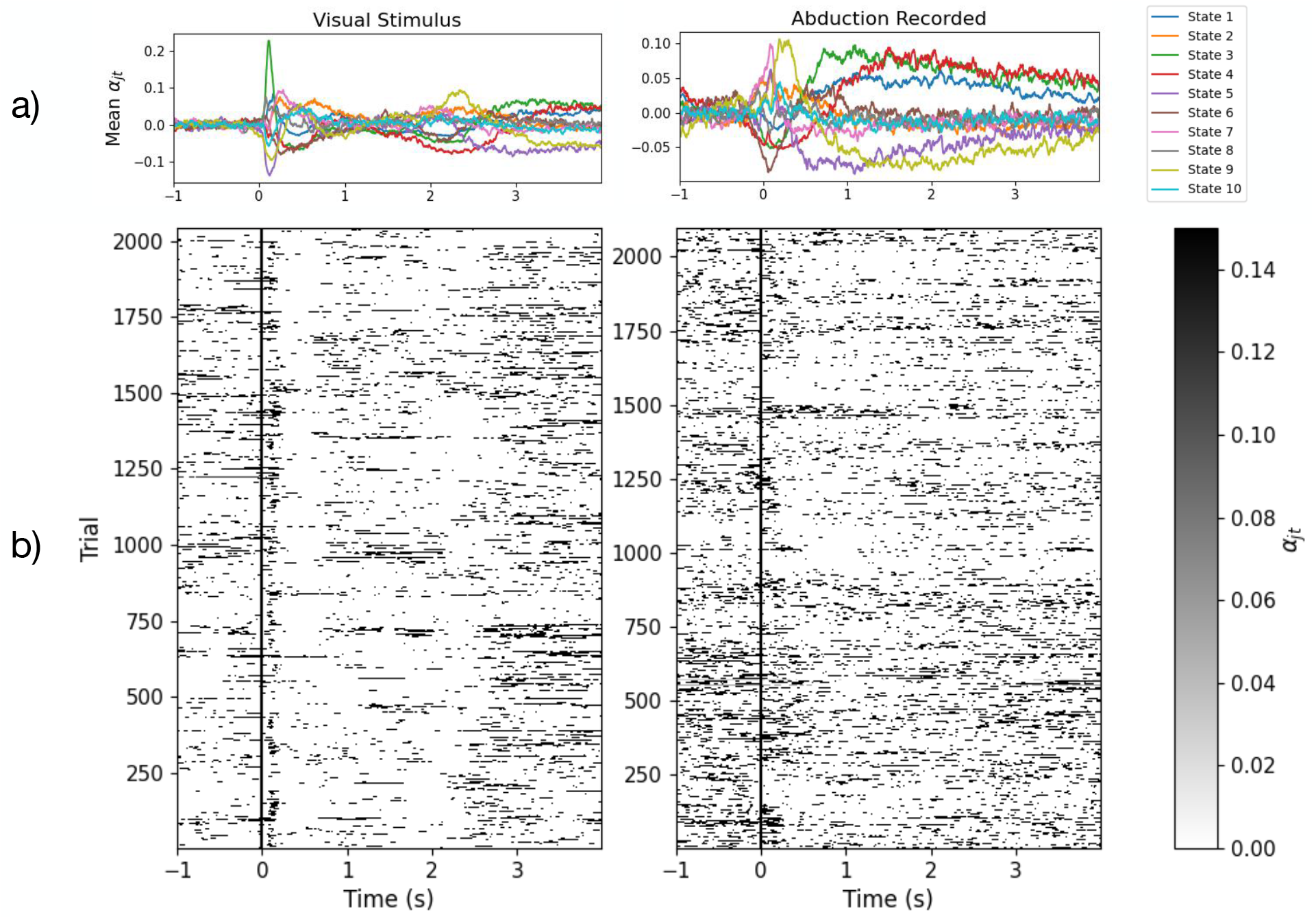
An evoked response to the visuomotor task is seen across trials with the HMM fit to the visuomotor task MEG dataset. a) State time courses epoched around the visual (left) and abduction (right) task. The average over trials is shown. b) Individual trial responses (state time course) for state 3 (visual, left) and state 5 (sensorimotor, right). The visual stimulus/abduction task occurs at Time = 0 s.

## Footnotes

1 Including the positivity constraint enables us to interpret the ***α***_*t*_ values as mixing coefficients and the sum to one constraint ensures the distribution of mixing coefficients is sufficiently non-Gaussian for the model to be identifiable [50].

2 Note, standardisation was also performed before PCA.

3 We only use the maximum a posteriori probability estimate post-hoc, during training we sample from the variational posterior distribution using the reparameterisation trick.

4 https://github.com/OHBA-analysis/HMM-MAR.

5 The Viterbi path is the maximum a posteriori probability estimate for the state in an HMM.

6 Also known as the dwell time.

7 logit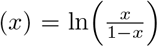.

8 It is possible to extract mode PSDs from the mode covariance matrices by reversing the PCA, which gives the auto-covariance of the time-delay embedded data. This matrix contains an estimate of the autocorrelation function, which can be Fourier transformed to give a mode PSD. However, the resolution of this PSD is limited by the number of time-delay embeddings, which results in a low-resolution PSD. This is why estimating the mode PSD using the inferred mixing coefficients and source reconstructed data is preferred.

